# Wnt/PCP-primed intestinal stem cells directly differentiate into enteroendocrine or Paneth cells

**DOI:** 10.1101/2020.08.12.247585

**Authors:** Anika Böttcher, Maren Büttner, Sophie Tritschler, Michael Sterr, Alexandra Aliluev, Lena Oppenländer, Ingo Burtscher, Steffen Sass, Martin Irmler, Johannes Beckers, Christoph Ziegenhain, Wolfgang Enard, Andrea C. Schamberger, Fien M. Verhamme, Oliver Eickelberg, Fabian J. Theis, Heiko Lickert

## Abstract

A detailed understanding of intestinal stem cell (ISC) self-renewal and differentiation is required to better treat chronic intestinal diseases. However, different models of ISC lineage hierarchy^1–6^ and segregation^7–12^ are debated. Here we report the identification of Lgr5^+^ ISCs that express *Flattop* (*Fltp*), a Wnt/planar cell polarity (PCP) reporter and effector gene. Lineage labelling revealed that Wnt/PCP-activated *Fltp*^+^ ISCs are primed either towards the enteroendocrine or the Paneth cell lineage *in vivo*. Integration of time-resolved lineage labelling with genome-wide and targeted single-cell gene expression analysis allowed us to delineate the ISC differentiation path into enteroendocrine and Paneth cells at the molecular level. Strikingly, we found that both lineages are directly recruited from ISCs via unipotent transition states, challenging the existence of formerly predicted bi- or multipotent secretory progenitors^7–12^. Transitory cells that mature into Paneth cells are quiescent and express both stem cell and secretory lineage genes, indicating that these cells are the previously described Lgr5^+^ label-retaining cells^7^. Wnt/PCP-activated Lgr5^+^ ISCs are indistinguishable from Wnt/β-catenin-activated Lgr5^+^ ISCs based on the expression of stem-cell signature or secretory lineagespecifying genes but possess less self-renewal activity. This suggests that lineage priming and cell-cycle exit is triggered at the post-transcriptional level by polarity cues and a switch from canonical to non-canonical Wnt/PCP signalling. Taken together, we identified the Wnt/PCP pathway as a new niche signal and polarity cue regulating stem cell fate. Active Wnt/PCP signalling represents one of the earliest events in ISC lineage priming towards the Paneth and enteroendocrine cell fate, preceding lateral inhibition and expression of secretory lineage-specifying genes. Thus, our findings provide a better understanding of the niche signals and redefine the mechanisms underlying ISC lineage hierarchy and segregation.

---

The intestinal epithelium renews continuously throughout life from a pool of Wnt/β-catenin-dependent Lgr5^+^ intestinal stem cells (ISCs)^1,13^. Lineage commitment of Lgr5^+^ ISCs has been viewed as a binary decision between an absorptive and a secretory progenitor through Notch/Delta-mediated lateral inhibition^14,15^. However, controversies exist regarding the potency of secretory progenitors and their differentiation routes into goblet cells, tuft cells, Paneth cells (PCs) or enteroendocrine cells (EECs)^8–12^. One possibility is that ISCs are functionally heterogeneous and directly differentiate into secretory cell types without passing through proposed stages of bi- or multipotent secretory progenitors. Indeed, functional heterogeneity of ISCs is evident in that i) ISCs positioned at the centre and the periphery of the crypt base produce different clone sizes^16^, ii) reserve stem cells are described at crypt position +4^17–20^ and iii) not all Lgr5^+^ cells constitute functional ISCs *in vivo*^21^ and some might correspond to non-cycling Lgr5^+^ label-retaining cells (LRCs), which are PC and/or EEC progenitors^7^. The non-canonical Wnt/planar cell polarity (PCP) pathway controls beta cell differentiation in the pancreas^22,23^ and determines functional heterogeneity in the islets of Langerhans^23,24^. Remarkably, the canonical Wnt/β-catenin pathway (which is required for the maintenance of Lgr5^+^ ISCs) and the non-canonical Wnt/PCP pathway share signalling components and antagonize each other’s function in some tissues^25,26^. Hence, Wnt/PCP signalling is a prime candidate pathway to control ISC heterogeneity and lineage choice.

## Fltp^+^ ISCs are fate biased

To analyse the impact of Wnt/PCP activation on ISC self-renewal and differentiation in the crypt stem cell niche we took advantage of the first reported sentinel for pathway activation, our knock-in *Fltp^ZV^* (Lac<u>Z</u> and H2B-<u>V</u>enus) dual reporter mouse model (Fig. 1a)^23,27^. In this mouse model, *Fltp*-H2B Venus reporter (FVR) activity is cell-cycle dependent and restricted to quiescent and terminally differentiated cells that had previously induced *Fltp* expression during Wnt/PCP acquisition^23,27^. When analysing small intestinal (SI) crypt cells in this model, we could distinguish three populations based on their FVR label intensity: FVR^neg^, FVR^low^ and FVR^hi^ cells (Fig. 1b). We isolated these populations using flow cytometry and characterized them using genome-wide transcriptional profiling (Fig. 1c and Extended Data Fig. 1a, b). KEGG pathway analysis revealed that the FVR^neg^, FVR^low^ and FVR^hi^ populations can be distinguished by their expression of genes involved in absorption, hormone secretion and immune-regulation, respectively, and by genes regulating cell-cycle behaviour and metabolic activity (Extended Data Fig. 1c and Supplementary Table 1). FVR^low^ and FVR^hi^ cells were highly enriched in transcripts associated with different EEC subsets (*Chga, Chgb, Nkx2-2, Neurod1*) and the PC lineage (*Nupr1, Dll4, Mmp7, Lyz1*), respectively, when compared to Lgr5^high^ ISCs (Fig. 1c and Extended Data Fig. 1d, e). Further, the FVR^hi^ population coexpressed ISC markers (Fig. 1c). This combined expression pattern is similar to the transcriptional profile of quiescent Lgr5^+^ LRCs (Fig. 1c)^7^. The low expression of cell proliferation genes (*Ccnd1, Ki67*) and the high level of the cell-cycle inhibitor gene *Cdkn1a* in both FVR^+^ populations suggested that FVR^+^ cells undergo terminal differentiation (Extended Data Fig. 1f). Indeed, FVR activity labelled a subset of the postmitotic secretory lineage namely essentially all Lyz1^+^ PCs (95.58%) and ChgA^+^ crypt EECs (96.13%), but not goblet or tuft cells (Fig. 1d-f and Extended Data Fig. 1g-i). Using a targeted single-cell qRT-PCR approach, we found that *Fltp* expression was restricted to a few FVR^low^ EECs and FVR^hi^ PCs, and to a subset of Lgr5^+^ ISCs (6.2%) (Fig. 1g). *Fltp* mRNA expression was induced by Wnt/PCP ligand stimulation indicating that Fltp is also a Wnt/PCP effector in the gut (Fig. 1h). From these data we conclude that i) *Fltp* is transiently expressed in ISCs, and ii) FVR due to reporter protein stability labels the immediate daughter cells of *Fltp*^+^ ISCs; i.e., more than 95% of the EECs and PCs.

**Figure 1.**
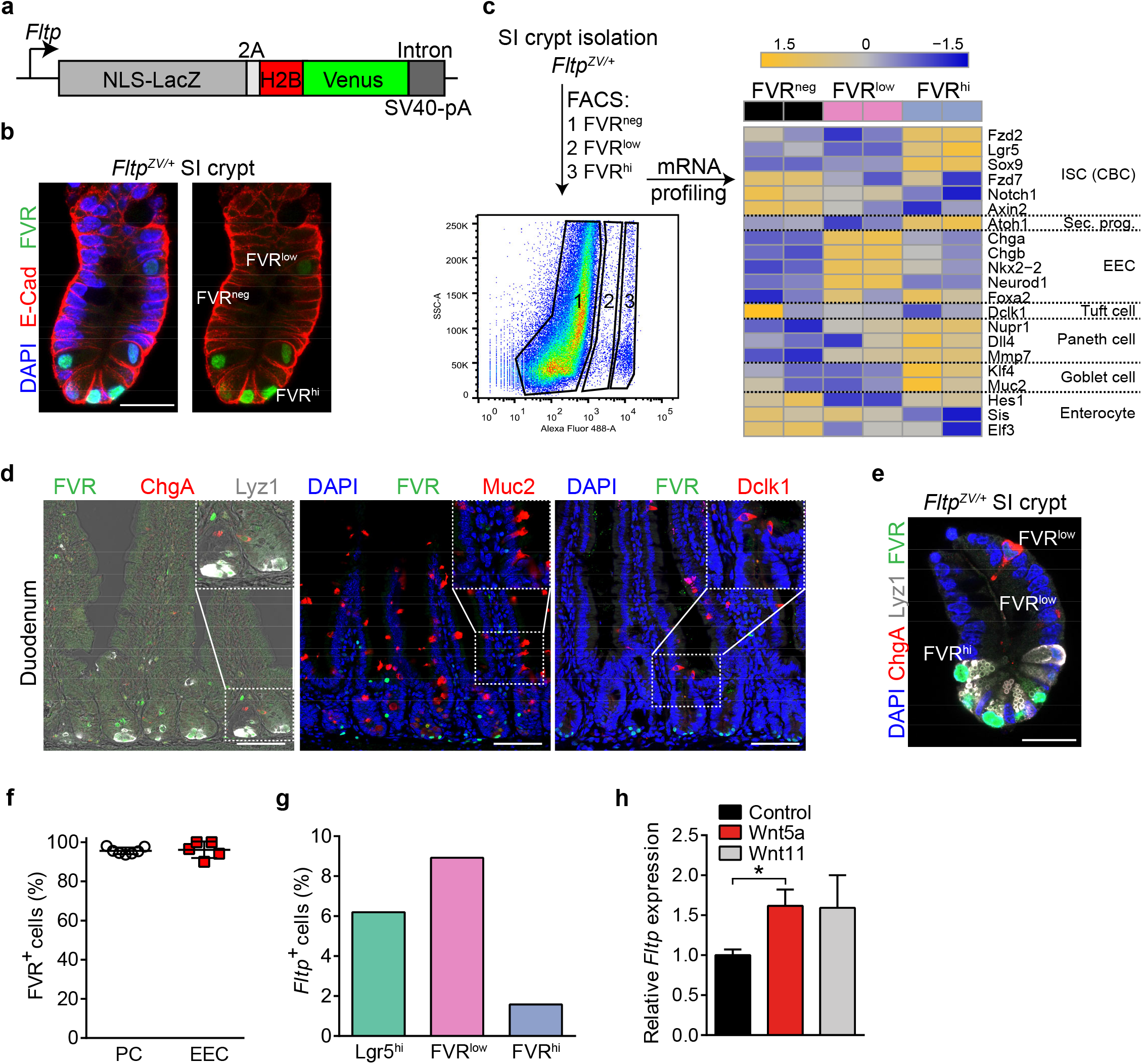
FVR labels the secretory enteroendocrine and Paneth cell lineages. **a,** Schematic of the loss-of-function and NLS-LacZ/H2B-Venus transcriptional reporter allele (*Fltp^ZV^*) for *Fltp*. **b,** Laser scanning confocal microscopy (LSM) image of a representative small intestinal (SI) crypt isolated from a *Fltp^ZV/+^* reporter mouse line depicting FVR^hi/low/neg^ crypt cells. FVR (Venus, green), DAPI (blue, nuclei), E-cad (red, membrane). Scale bar, 25 μm. **c,** Experimental design for microarray analysis of the three small intestinal (SI) crypt cell populations distinguishable by FVR activity. The heatmap depicts the expression profiles of key stem-cell and intestinal lineage genes. Expression is scaled row-wise and the colours range from dark blue (low expression) to orange (high expression) and represent normalized expression (row z-score). ISC, intestinal stem cell. CBC, crypt base columnar cell. Sec. progenitor, secretory progenitor. EEC, enteroendocrine cell. **d,** LSM images showing FVR (Venus, green) expression in the secretory lineages in the adult intestine of *Fltp^ZV/+^* mice (4 months) co-stained against ChgA (red, enteroendocrine cells), Lyz1 (white, Paneth cells), Muc2 (red, goblet cells), and Dclk1 (red, tuft cells). DAPI (blue) stains the nucleus. Scale bars, 75 μm. **e, f,** LSM image depicting a representative SI crypt isolated from adult *Fltp^ZV/+^* reporter mice with indicated FVR^neg/low/hi^ cells stained for DAPI (blue, nucleus), FVR (Venus, green), ChgA (red, enteroendocrine cells), and Lyz1 (white, Paneth cells) (**e**) and quantification of Lyz1^+^ FVR^+^ Paneth cells (PCs) and ChgA^+^ FVR^+^ enteroendocrine cells (EECs) (**f**). For Paneth cells: *n* (mice) = 8 with 99 analysed crypts (95.58 % of PCs = FVR^+^). For enteroendocrine cells: *n* (mice) = 5 with 76 analysed crypts (96.13 % of EECs = FVR^+^). Scale bar, 25 μm. **g,** Relative abundance of *Fltp^+^* Lgr5^hi^, FVR^low^ and FVR^hi^ cells determined by single-cell qRT-PCR. *n* = 145 Lgr5^hi^ cells; *n* = 112 FVR^low^ cells; *n* = 126 FVR^hi^ cells. *n* (mice) = 3 for Fltp^ZV/+^, Lgr5-ki **h,** *Fltp* expression in crypts treated with indicated Wnt/PCP ligands for two days. *n* (independent experiments) = 4. Error bars represent SEM. Two-tailed Student’s *t*-test, \**P*<0.05, \*\*\**P*<0.001.

## Wnt/PCP activated Fltp^+^ ISCs and Wnt/β-catenin activated Lgr5^+^ ISCs are indistinguishable at the transcriptional level

Homogeneity and equipotency of the ISC pool is vividly discussed^3–5,16–19,21,28,29^. Our data suggested that *Fltp*^+^ ISCs are primed towards the EEC and/or PC lineage. To determine the mechanism underlying ISC heterogeneity and lineage priming, we next used a dual-fluorescent Cre-reporter *Fltp^T2AiCre/+^; Gt(ROSA)26^mTmG/+^* mouse line (Extended Data Fig. 2a)^30,31^. In this model, Cre recombinase induces a switch from membrane-Tomato (mT) to membrane-GFP (mG). The intermediate mTmG state can be captured by flow cytometry and highly expresses *Fltp* (Fig. 2a-c). *In vivo, Fltp^+^* mTmG cells predominantly located at crypt position +4/+5 (Fig. 2d, e). Consistent with the transient expression of the Wnt/PCP reporter gene *Fltp* in a subset of Lgr5^+^ ISCs (Fig. 1g), the Wnt/PCP activated *Fltp^+^* mTmG cells and Wnt/β-catenin activated Lgr5^+^ ISCs are closely related as exemplified by similar expression of the Lgr5^+^ ISC signature genes^28^, lineage-specifying genes and Notch pathway genes, as well as the lack of Lgr5^+^ LRC markers^7^ (Fig. 2f-i and Extended Data Fig. 2b, c). However, we found that less mTmG cells were in cell-cycle and formed organoids compared to Lgr5^+^ ISCs (Extended Data Fig. 2d-i). As *Fltp*^+^ mTmG cells possessed limited self-renewal capacity *in vitro* we next analysed whether chemical injury of the intestine can activate these cells. In contrast to Wnt/β-catenin activated Lgr5^+^ ISCs, mTmG cells were resistant to 5-FU treatment but showed reduced mitotic activity (Extended Data Fig. 3a-i). We sporadically detected lineage-traced villi in the small intestine in homeostasis and the number did not increase in 5-FU treated mice indicating that mTmG cells do not contribute to regeneration after intestinal injury (Extended Data Fig. 3j-l). Together, these data show that Fltp^+^ ISCs are transcriptionally similar to Lgr5^+^ ISCs but possess limited self-renewal capacity *in vitro* and *in vivo*. Further, we conclude that Wnt/PCP signalling-induced lineage priming in Fltp^+^ ISCs represents the earliest step in the commitment of ISCs towards the PC or EEC lineage. These early cell fate decisions are triggered at the post-transcriptional level by polarity cues and precede Notch/Delta-mediated lateral inhibition.

**Figure 2.**
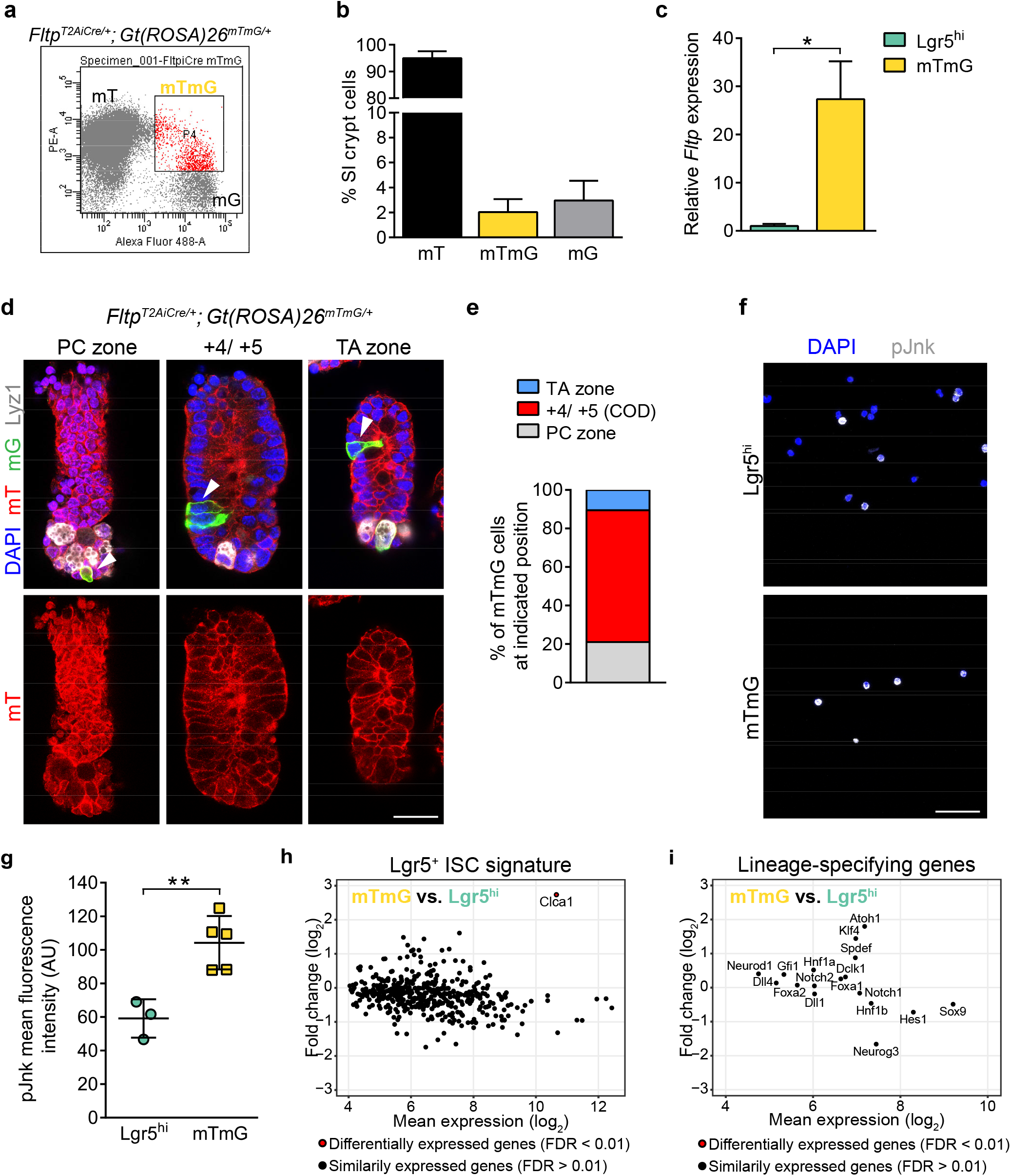
Wnt/β-catenin and Wnt/PCP activated Lgr5^+^ ISCs are indistinguishable at the transcriptional level. **a, b**, Flow cytometry analysis (**a**) and relative abundance (**b**) of Fltp lineage^-^ (mT), Fltp^+^ intermediate cells (mTmG) and Fltp lineage^+^ (mG) crypt cells isolated from *Fltp^T2AiCre/+^; Gt(ROSA)26^mTmG/+^* mice. *n* (mice) = 6. Error bars represent SD. **c,** qRT-PCR data comparing relative *Fltp* expression in mTmG crypt cells and Lgr5^hi^ intestinal stem cells. *n* (independent experiments) = 3 for Lgr5^hi^ cells. *n* (independent experiments) = 2 (4 mice) for mTmG cells. Error bars represent SD. Two-tailed Students’s *t*-test, \**P*<0.05. **d, e,** Representative LSM images of Fltp^+^ intermediate mTmG cells at indicated positions. SI crypts from *Fltp^T2AiCre/+^;Gt(ROSA)26^mTmG/+^* mice were stained for DAPI (blue, nucleus), mT (Fltp lineage^-^, RFP, green), mG (Fltp^+^ intermediate cells, green, GFP), and Lyz1 (white, Paneth cells). Scale bar, 25 μm. (**d**). Abundance of mTmG cells at indicated positions (**e**). *n* (mice) = 3. **f, g,** Representative LSM images of Lgr5^hi^ and mTmG cells isolated by flow cytometry and stained for active, phosphorylated Jun N-terminal kinase (pJnk, white) indicating active Wnt/PCP signalling and DAPI (blue, stains nuclei) (**f**). Quantification of the mean fluorescent intensity of pJnk (**g**). *n* (independent experiments) = 3 for Lgr5^hi^ cells. *n* (independent experiments) = 5 for mTmG cells. Error bars represent SD. Two-tailed Student’s *t*-test, \**P*<0.05. Scale bar, 50 μm. **h,** MA-plot comparing the expression of ISC signature genes in mTmG and Lgr5^hi^ (ISCs) cells. Differentially expressed genes (FDR < 0.01) are indicated in red. The y-axis indicates the fold change in log2 and the x-axis indicates the mean log2 expression value. *Clca1* is the only significantly regulated gene. *n* (Lgr5^hi^ ISC microarray samples) = 6. *n* (mTmG microarray samples) = 4. **i,** MA-plot comparing the expression of lineage-specifying genes in mTmG and Lgr5^hi^ cells. The y-axis indicates the fold change in log2 and the x-axis indicates the mean log2 expression value. No gene is significantly regulated. *n* (microarray samples) = 6 for Lgr5^hi^ ISC. *n* (microarray samples) = 4 for mTmG.

## Intestinal lineages form directly from ISCs via unipotent lineage-specified progenitors

Secretory lineage specification and in particular the signalling pathways that regulate secretory subtype specification remain poorly understood. Multi-potent secretory progenitors that specify during multiple rounds of division^11,15^, in addition to different lineage-restricted bi-potent secretory progenitors^7,9,10,12^ and direct differentiation from ISCs^2^, have been proposed (Extended Data Fig. 4a, b). Our data suggested that the PC and EEC lineage directly allocate from ISCs. To elucidate the lineage hierarchy and conclude on lineage relationships in the intestinal epithelium, we made use of the *Fltp^ZV^* and Foxa2 Venus Fusion (FVF) reporter mouse lines^27,32^ that label rare intestinal cell populations and performed scRNAseq of 60,000 cells in homeostasis (Fig. 3a, Supplementary Table 2). Using this approach, we could highly enrich for the PC, EEC, goblet and tuft cell lineages (Fig. 3b, c, Extended Data Fig. 4c, d and Supplementary Table 3). Strikingly, due to the enrichment and high resolution of the secretory lineages we could identify all described intestinal epithelial populations and in addition lineage-specified progenitors with proliferative activity and distinct expression profiles (Fig. 3b-e, Extended Data Fig. 4c-g, Extended Data Fig. 5a-c, Supplementary Table 4). Further, to computational reconstruct possible lineage relationships and differentiation trajectories we used partitioned graph abstraction (PAGA)^33^. Strikingly, we found that all lineages originated from ISCs. Progenitor states were highly interconnected reflecting the high plasticity of intestinal epithelial cells^34^ (Extended Data Fig. 5d). Together, these data imply that for every intestinal lineage a progenitor with a distinct transcriptional signature exists and that all lineages directly allocate from ISCs.

**Figure 3.**
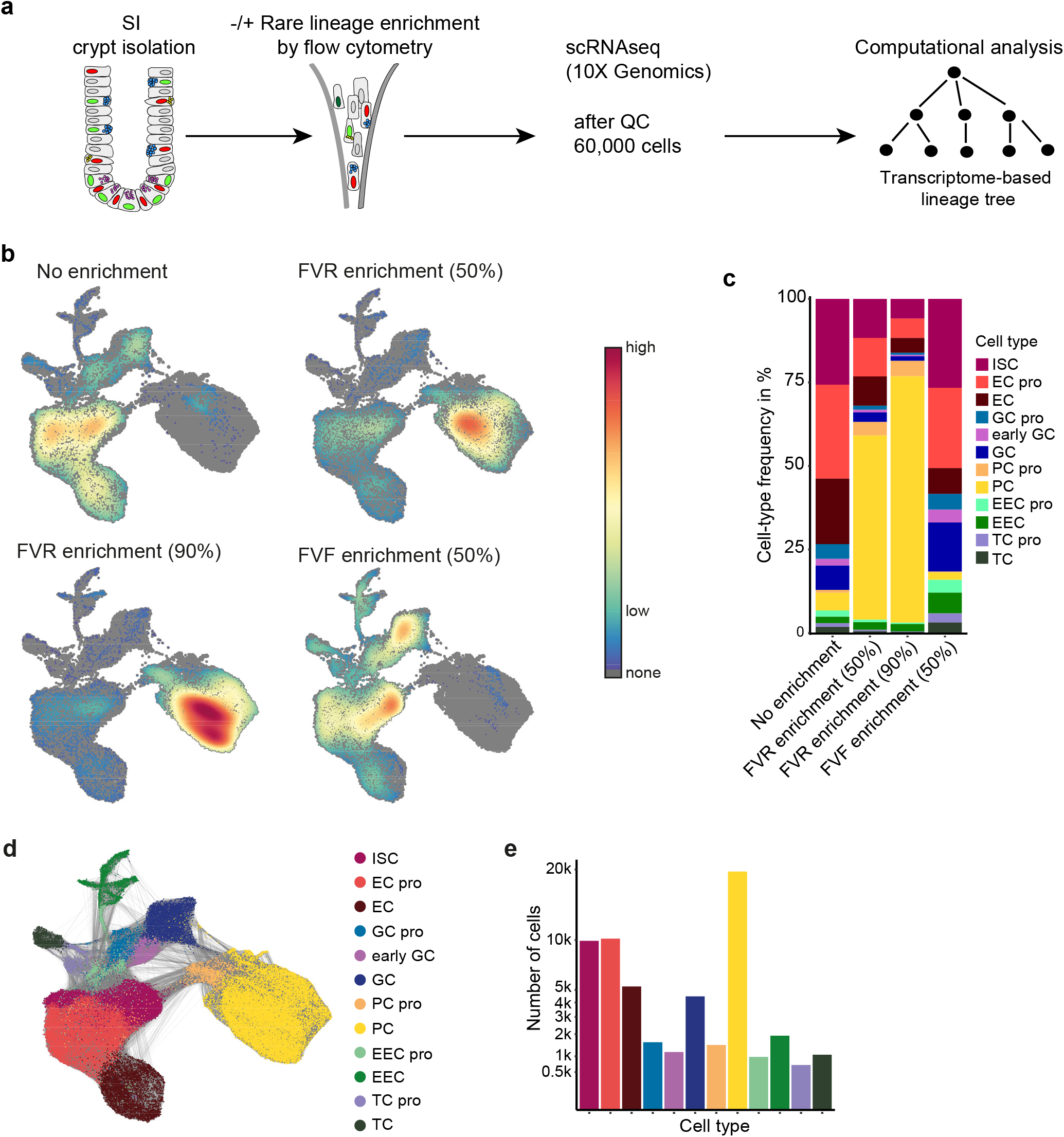
Subtype enrichment of crypt cells reveals distinct progenitor states for all intestinal lineages. **a,** Experimental overview. SI crypt cells were obtained from reporter (FVR, FVF) mice to enrich rare cell states, and wild-type mice. Fluorescence-activated cell sorting (FACS) was used to enrich rare crypt cell populations from reporter mice. Transcriptional profiling of single cells was performed using the 10X Genomics platform. **b,** Reporter mice enable the identification of rare cell states. Cell-density plots of FVR and FVF based enrichment compared to a non-enriched sample. No enrichment (pooled cells from 2 mice), FVR enrichment 50% (pooled cells from 2 mice, flow-enriched FVR^+^ cells were mixed with nonenriched cells in a ratio of 1:2), FVR enrichment 90% (pooled cells from 2 mice, flow-enriched FVR^+^ cells were mixed with non-enriched cells in a ratio of 9:1), FVV enrichment 50% (n (mice) = 3, flow-enriched FVF^+^ cells were mixed with non-enriched cells in a ratio of 1:2. **c,** Bar plot depicting the relative abundance of progenitor and mature cell types in the nonenriched, and FVR and FVF enriched sample. FVR enrichment enables the isolation of rare PC progenitors and mature PCs whereas FVF enrichment enables the efficient extraction of the EEC and goblet cell lineage compared to the non-enriched sample. ISC, intestinal stem cell. EC pro, enterocyte progenitor. EC, enterocyte. GC pro, goblet cell progenitor. Early GC, early goblet cell. GC, mature goblet cell. PC pro, Paneth cell progenitor. PC, Paneth cell. EEC pro, enteroendocrine cell progenitor. EEC, enteroendocrine cell. TC pro, tuft cell progenitor. TC, tuft cell. **d,** UMAP plots of all control intestinal crypt cells highlighting progenitor cell-type annotation. Grey lines depict 30 nearest neighbors for each cell. Cells were obtained from 10 samples including wild-type (control) and reporter mice for lineage enrichment (Supplementary Table 2). **e,** The bar plot depicts the number of captured cells per cell type.

## Integration of temporal-resolved lineage labelling with single-cell expression analysis confirms unipotent secretory progenitors

As transcriptomes alone may not accurately determine the future cell fate of ISCs and progenitors^35,36^ we combined temporal-resolved lineage labelling with a highly sensitive singlecell qPCR approach to fine-map fate decisions towards the PC and EEC lineage (Fig. 4a, b). We determined the expression of 80 well-known and functionally important intestinal signature genes from defined molecular categories; i.e., stem-cell signature genes, lineage markers and determinants, signalling components and cell-cycle regulators (Fig. 4b and Supplementary Table 5). To gain temporal and progenitor state resolution we included: i) Lgr5^hi^ ISCs from *Lgr5-ki* mice, ii) early and late mTmG cells from *Fltp^T2AiCre/+^; Gt(ROSA)26^mTmG/+^* mice, iii) FVR^+^/Lgr5^hi^ and FVR^+^/Lgr5^low^ double-positive cells from *Fltp^ZV^/Lgr5-ki* dual reporter mice (Fig. 4c-e), and iv) Neurog3-expressing EEC progenitors from Ngn3-Venus mice^37^. UMAP visualization of the single-cell qPCR data showed that the flow-sorted FVR^hi^ PCs grouped into one defined cluster, whereas the FVR^low^ EECs grouped into two clusters (Fig. 4f). Early mTmG cells grouped together with Lgr5^hi^ ISCs and late mTmG cells grouped together with a subset of FVR^low^ cells (Fig. 4f).

**Figure 4.**
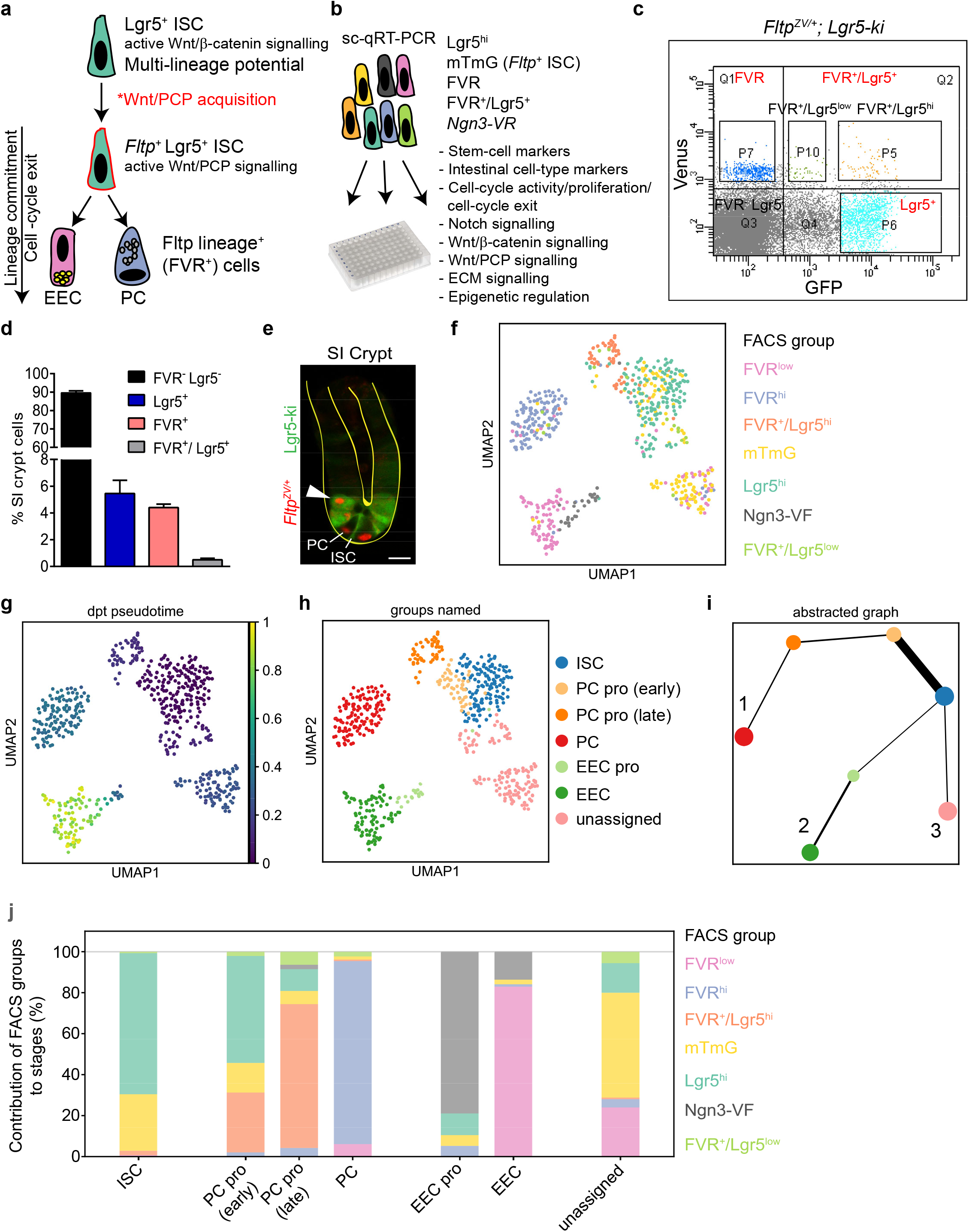
Temporal-resolved lineage labelling and pseudotemporal ordering of intestinal crypt cells shows that EECs and PCs directly allocate from ISCs via unipotent transition states. **a, b,** Wnt/PCP activated *Fltp^+^* ISCs are committed to differentiate into Paneth cells (PCs) and enteroendocrine cells (EECs) (**a**). Integration of lineage labelling and single-cell gene expression to elucidate the differentiation trajectories from ISCs into the Paneth and enteroendocrine lineage by single-cell qRT-PCR analysis of 80 genes from defined categories in Lgr5^hi^ ISCs, mTmG cells, FVR cells, FVR^+^/Lgr5^+^ cells and Ngn3-VR cells (**b**). **c, d,** FACS plot of crypt cells from *Fltp^ZV/+^; Lgr5-ki* dual reporter mice depicting the separation of rare FVR^+^/Lgr5^+^ (FVR^+^/Lgr5^hi^, FVR^+^/Lgr5^low^) cells from Lgr5^+^-GFP and FVR single positive, and Lgr5-GFP and FVR negative cells (**c**) and quantification of FACS analysis (**d**). 0.5% of the crypt cells are FVR^+^Lgr5^+^ double positive. *n* (independent experiments, mice) = 7. Error bars represent SEM. **e,** LSM live image of a representative SI crypt, cultured in matrigel, isolated from *Fltp^ZV/+^*; Lgr5-ki mice showing rare FVR (red), Lgr5 (green) double positive cells (arrowhead) located at position +4 (= supra-Paneth cell position). PC, Paneth cell. ISC, intestinal stem cell (Lgr5^+^). Scale bar, 10 μm. **f, g, h,** UMAP projection of Lgr5^hi^ cells, early and late mTmG cells, FVR^+^/Lgr5^+^ cells, FVR cells and Ngn3-VF endocrine progenitors based on the expression of 80 marker genes. Each dot represents a single cell. Colours indicate FACS groups (**f**), pseudotime computed using dpt (**g**), and cell-type clusters annotated based on marker genes (**h**). **i,** PAGA plot showing relationship of cell type clusters in (h). **j,** The bar plot depicts the contribution of FACS groups to stages in the PC and EEC branch.

To delineate the ISC differentiation path into the EEC and PC lineages we used PAGA^33^. Ordering of cells along a pseudotime as a proxy for real-time differentiation identified three terminal states from our single-cell qPCR snapshot data: the PC branch with terminal state 1, the EEC branch with terminal state 2 and a third branch with unassigned cells (Fig. 4g-j and Extended Data Fig. 6a). Separation into the EEC and PC lineages occurred early and still within the ISC population, which reinforces that EECs and PCs directly allocate from Lgr5^+^ ISCs (Fig. 4i, j and Extended Data Fig. 6b). Plotting gene expression versus pseudotime revealed that cells differentiate via lineage-specific unipotent transition states characterized by downregulation of stem cell markers for the EEC branch and co-expression of stem-cell and secretory markers for the PC branch (Extended Data Fig. 6b-f). Unassigned cells were mainly late mTmG and FVR^low^ cells (Fig. 4j) and did not express mature EEC or PC markers (Extended Data Fig. 6a), but instead were characterized by the expression of genes implicated in cell-matrix adhesion and signalling (*Itga1, ItgaV*), repression of endocrine cell fate (*Gfi1*) and regulation of Notch signalling (*Lfng*) (Extended Data Fig. 6b). Together, the pseudotemporal analysis further supports our realtime lineage reporter-based finding that PCs and EECs directly allocate from ISCs via unipotent progenitors.

## FVR^+^ LRCs differentiate into PCs

Single-cell transcriptomics data suggest that PCs are mainly formed via a PC/goblet cell precursor and that only a small subset directly differentiates from ISCs^2^. However, our integrated analysis of lineage labelling with single-cell gene expression indicates that in addition to EECs most of the PCs also directly allocate from ISCs via a unipotent transition state comprising mainly Lgr5^+^/FVR^+^ double-positive cells (Fig. 4j and Extended Data Fig. 6).

Using the FVR mice we obtained transcriptional profiles of more than 20,000 cells from the rare and difficult to capture PC lineage^29^ (Fig. 3). When we investigated the transition of ISCs to PCs we identified a ISC population that connects to the PC progenitor population, which we termed PC-primed ISCs (Fig. 5a). In addition, we found two mature PC types which differ in the expression of *Lyz2* (Fig. 5b). Pseudotemporal ordering of ISCs and PC subclusters placed the PC progenitor in between the mature PCs and ISCs, while PC-primed ISCs link to PC progenitors (Fig. 5c). The co-expression of stem-cell and secretory lineages genes in the PC progenitor suggested that these are the recently described quiescent Lgr5^+^ LRCs^7^ (Fig. 4, 5c, d and Extended Data Fig. 6a-c). With the exception of PCs that have a lifespan of about 6-8 weeks, all other differentiated intestinal lineages are renewed every 4-5 days. Quiescent or slowly cycling intestinal cells persist for more than 10 days and thus are defined by the property of label-retention. To assess label-retention within the FVR^+^ population we “birth-dated” this population with 5-bromodeoxyuridine (BrdU) (Fig. 5e-g). After a chase period of 10 days, 30% of the FVR^+^ cells retained the label and were non-PCs (Lyz1^-^), and hence LRCs (Fig. 5h). After a chase period of 21 days, all BrdU^+^ LRCs present in the crypt were also FVR^+^. An increase in FVR^+^/BrdU^+^/Lyz1^+^ cells implied that FVR^+^ LRCs primarily give rise to PCs (Fig. 5h). Like *Fltp^+^* ISCs (mTmG cells), FVR^+^ LRCs were predominantly located at position +4/+5 (Fig. 2d, e and Fig. 5i-k). These results indicate that PCs directly arise from ISCs via a quiescent, label-retaining transition state, which is characterized by co-expression of stem cell and secretory lineage markers. The identification of lineage-specific unipotent transition states together with the fact that FVR labels more than 95% of all PCs and EECs, but not goblet or tuft cells challenges the existence of formerly predicted bi- or multipotent secretory progenitors.

**Figure 5.**
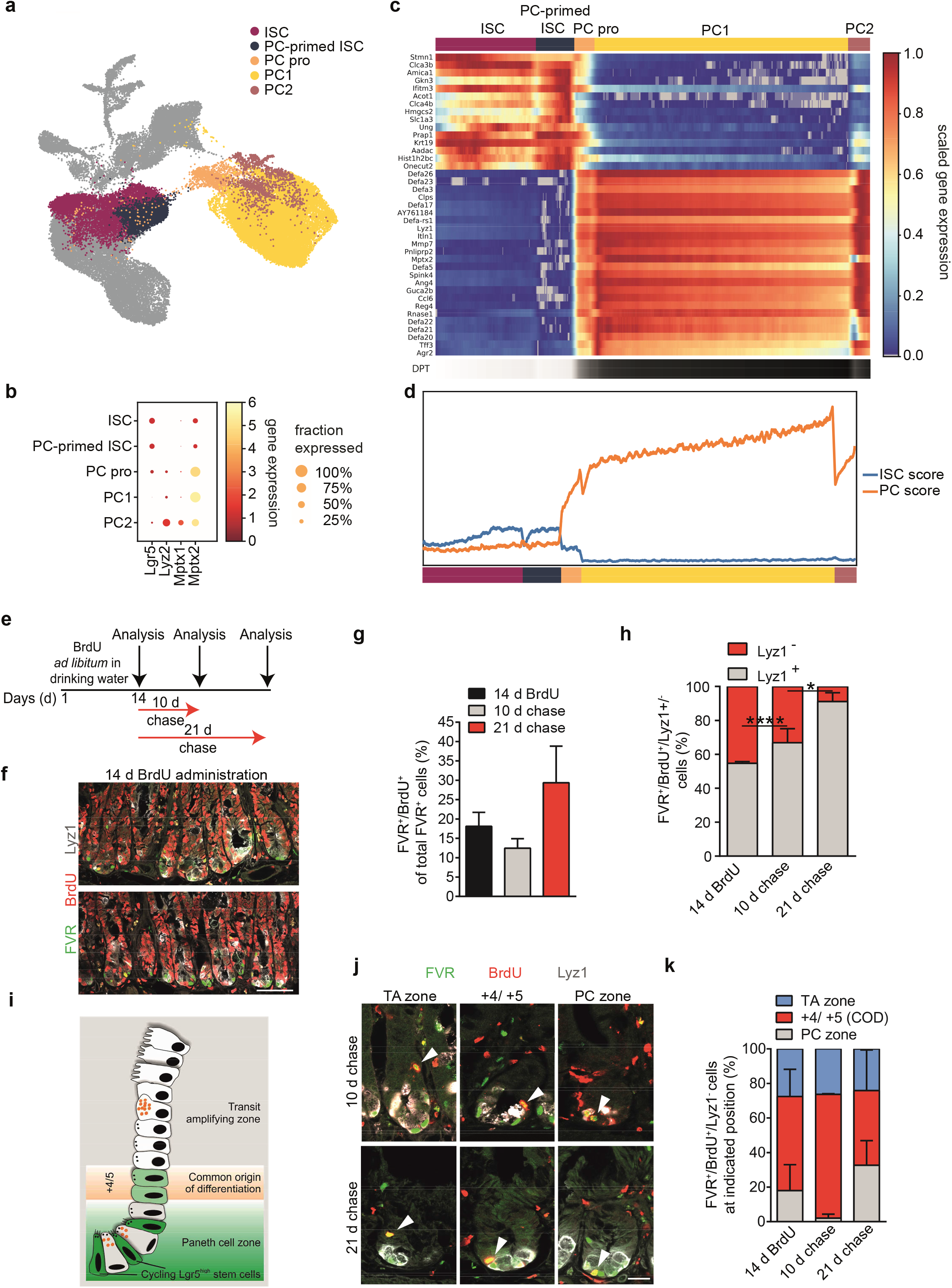
FVR marks Lgr5^+^ label-retaining cells that give rise to Paneth cells. **a,** UMAP plot depicting the identified cell states during differentiation of ISCs towards the Paneth cell lineage. Our analysis revealed an ISC subpopulation that is primed towards PC fate (PC-primed ISC), a PC progenitor population (PC pro), a PC population characterized by absence of *Lyz2* expression (PC1) and a PC population characterized by *Lyz2* expression (PC2). **b,** Dotplot showing the expression of the stem-cell marker *Lgr5* in PC progenitor populations and genes that distinguish the mature PC types PC1 and PC2. **c,** Smoothed gene expression heatmap of differentially expressed genes of the displayed cell states plotted along diffusion pseudotime from ISCs to mature PCs. **d,** ISC score and PC score along pseudotime shows high ISC score in PC-primed ISCs. Also, Paneth cell progenitors express both ISC and PC markers. **e,** Experimental scheme of the 5-bromo-2’-deoxyuridine (BrdU) pulse-chase experiment. To assess label-retention, *Fltp^ZV/+^* mice were treated with BrdU for 14 days (d) and analysed after a chase period of 10 and 21 d. **f,** Representative LSM images of duodenal sections stained for FVR (Venus, green), BrdU (red) and Lyz1 (white, Paneth cells) after 14 d BrdU administration. Scale bar, 25μm. **g,** Relative proportion of FVR^+^/BrdU^+^ cells of total FVR^+^ cells at the indicated time points. 14 d BrdU: *n* (mice) = 4 with 1432 analysed FVR^+^ cells; 10 d chase: *n* (mice) = 2 with 785 analysed FVR^+^ cells; 21 d chase: *n* (mice) = 4 with 2401 analysed FVR^+^ cells. Error bars represent SD. **h,** Relative proportion of Lyz1^+^ (long-living and newly formed Paneth cells) and Lyz1^-^ (LRC) cells of the total FVR^+^/BrdU^+^ cells at the indicated time points. 14 d BrdU: *n* (mice) = 4 with 1432 analysed FVR^+^ cells; 10 d chase: *n* (mice) = 2 with 785 analysed FVR^+^ cells; 21 d chase: *n* (mice) = 4 with 2401 analysed FVR^+^ cells. Error bars represent SD. Two-tailed Student’s *t*-test, \**P*<0.05, \*\*\*\**P*<0.0001. **i, j, k,** Scheme depicting the compartmentalization of the SI crypt. Actively cycling stem cells (Lgr5^+^) and Paneth cells reside in the Paneth cell (PC) zone. Quiescent/label-retaining cells locate at position +4/ +5. The transit-amplifying zone (TA zone) contains mainly proliferative (absorptive) progenitors (**i**). Representative LSM images from duodenal sections stained for FVR (Venus, green), BrdU (red) and Lyz1 (white, Paneth cells) after 10 d and 21 d chase. The position (arrowhead) of the FVR^+^ LRC (BrdU^+^/Lyz1^-^) is defined according to the scheme in (**i**). Quantification of FVR^+^ LRCs (BrdU^+^/Lyz1^-^) at indicated positions (according to j). 14 d BrdU: *n* (cells) = 116 (in 4 mice), 10 d chase: *n* (cells) = 32 (in 2 mice), 21 d chase: *n* (cells) = 55 (in 4 mice) (**k**). Scale bar, 25 μm (**j**). Error bars represent SD.

## Wnt/PCP signalling is activated during PC and EEC differentiation

Fltp is transiently expressed in ISCs that acquired Wnt/PCP activation and are committed to differentiate into PCs and EECs (Fig. 1g, Fig. 2f, g). Cell ordering along the pseudotime confirmed that i) Wnt/PCP signalling is specifically activated in ISCs that differentiate towards PCs and EECs indicated by upregulation of several Wnt/PCP genes such as *Vangl2, Dvl2, Ror2* and *Celsr1* and ii) Wnt/PCP pathway activation precedes Notch/Delta-mediated lateral inhibition and cell-cycle exit (Extended Data Fig. 7a, b and Extended Data Fig. 2b). Consistent with the expression of core pathway components we also detected increased expression of Wnt/PCP pathway genes and Jnk activity in the FVR^+^ population, indicative of active Wnt/PCP signalling (Extended Data Fig. 7c-f). To further corroborate a role of Wnt/PCP signalling in cell fate regulation we analysed Celsr1^crsh/+^; *Fltp^ZV/ZV^* compound mutant mice. Celsr1 is a core Wnt/PCP component and member of a family consisting of Celsr1-3. Gene expression analysis of 14,000 mutant cells (from *n=4* mutant mice) revealed a less proliferative PC progenitor population and aberrant gene expression of known secretory lineage regulators (e.g. *Sox9, Atoh1, Spdef Foxa3, Tead2* and *Jun*) and potentially new regulators (e.g. *Ybx1, Pa2g4, Hnrnpk*) as well as canonical Wnt target genes specifically in the PC lineage suggesting disturbances in the differentiation of PCs (Extended Data Fig. 8a-e). Assessment of PC and EEC numbers showed a slight reduction of PCs when compared to control mice (Extended Data Fig. 8f, g). These weak disturbances in gene expression and PC and EEC numbers in this mutant mouse model is most likely due to functional redundancy of Wnt/PCP proteins during PC and EEC differentiation (Extended Data Fig. 7).

Taken together, we identified the Wnt/PCP pathway as a new niche signal that determines stem cell fate. We propose that a switch from Wnt/β-catenin to Wnt/PCP signalling induces PC and EEC lineage priming and cell-cycle exit of ISCs. This is consistent with our recent findings that Wnt/PCP activation triggers functional maturation and cell-cycle exit of endocrine insulin-producing β-cells in the pancreatic islet^23^ and suggests that polarity cues regulate cell heterogeneity and terminal differentiation in the crypt and islet cell niche.

## Methods and Supplementary material

### Animal studies

Animal experiments were carried out in compliance with the German Animal Protection Act and with the approved guidelines of the Society of Laboratory Animals (GV-SOLAS) and of the Federation of Laboratory Animal Science Associations (FELASA). This study was approved by the institutional Animal Welfare Officer and by the Government of Upper Bavaria, Germany.

Mouse lines used: *Fltp^ZV^* (C57BL/6J)^27^, *Fltp^T2AiCre^* (mixed C57BL/6J, CD1 background)^30^ crossed with *Gt(ROSA)26^mTmG^* (mixed 129/SvJ, C57BL/6J background)^31^, *Lgr5-EGFP-IRES-creERT2* (C57BL/6J)^1^, Celsr1^Crsh^ (mixed BALB/c, C57BL/6J background)^38^, Ngn3-VF (mixed 129/SvJ, C57BL/6J background)^37^, homozygous Foxa2-Venus fusion (FVF)^39^ mice were generated as previously described and backcrossed to C57BL/6 background for at least 10 generations. All experiments were performed using 3-6-month-old mice, unless indicated otherwise.

### Tissue preparation and immunohistochemistry

The intestine was isolated and flushed with ice-cold PBS, fixed with 4% paraformaldehyde (PFA) for 3h at 4°C and then placed for cryoprotection in a progressive sucrose gradient (7.5% sucrose for 1h, 15% sucrose for 1h, 30% sucrose overnight). Tissue was embedded in Optimum Cutting Temperature (Leica Biosystems, Germany, #14020108926) and sectioned at 14μm. Isolated intestinal crypts were fixed for whole-mount stainings with 4% PFA for 30min at room temperature (RT). After fixation crypts were washed three times with PBS. For immunofluorescence staining, sections or isolated crypts were permeabilized with 0.5 % Triton X-100 in PBS for 30 min at RT, blocked [(10% FCS, 0.1% BSA and 3% donkey serum in PBS/0.1 %Tween-20 (PBST)] for 1 h and incubated with primary antibodies overnight at 4 °C. Sections or crypts were washed in PBST, incubated with secondary antibodies in blocking solution for 1h at RT, followed by a DAPI (ROTH, 6335.1) staining to visualize the nuclei and mounted with the Elvanol antifade reagent.

For BrdU staining tissue was sectioned at 14μm and stained according to standard procedure followed by incubation in 3.3N HCl for 10min on ice, 50min at 37°C and incubation with borate buffer pH8.5 for 2x 15min at RT.

For stainings on single cells (Ki67, pJnk, EdU, Venus) cells were isolated by flow cytometry and cytospun on glass slides. Cells were dried and fixed with 4% PFA for 10min at RT, permeabilized with 0.25% Triton-X100 in PBS for 15min at RT and then blocked for 1h at RT followed by an overnight incubation with the primary antibody. Cells were washed in PBST, incubated with secondary antibodies in blocking solution for 1h at RT, followed by a DAPI (ROTH, 6335.1) staining to visualize the nuclei and mounted with the Elvanol antifade reagent. Sections, cells and crypts were visualized using a Leica SP5 confocal microscope.

The following primary antibodies were used for immunofluorescent stainings: chicken anti-GFP (1:600, Aves Labs, USA, GFP-1020); rat anti-BrdU (1:200, Abcam, ab6326); rat anti-RFP (1:500, Chromotek, ORD003515), goat anti-ChgA (1:200, Santa Cruz, sc-1488); rabbit anti-Lyz1 (1:1000, DAKO, A0099); rabbit anti-Muc2 (1:500, Santa Cruz, sc-7314); rabbit anti-Dclk1 (1:200, Abcam, ab37994); rat anti-BrdU (1:200, Abcam, ab6326); rabbit anti-Ki67 (1:200, Abcam, ab15580); rabbit anti-pJnk (1:100, NEB, #4668). The following secondary antibodies were used: donkey anti-chicken Alexa Fluor 488 (Dianova, 703-225-155); donkey anti-mouse Cy5 (Dianova, 715-175-151); donkey anti-goat Alexa Fluor 555 (Invitrogen, A21432); donkey anti-rabbit Alexa Fluor 555 (Invitrogen, A31572); donkey anti-rabbit Alexa Fluor 649 (Dianova, 711-605-152); donkey anti-rat DyLight 549 (Dianova, #712-505-153).

### pJnk fluorescence intensity analysis

Lgr5^hi^ and mTmG cells were isolated by flow cytometry and cytospun on glass slides. Cells were stained and imaged using a Leica SP5 Confocal microscope. Analysis was performed using the Leica LAS-AF (Version 2.7.3.9723) software. pJnk fluorescent intensity signal was determined from each cell and background signal was subtracted (secondary antibody only).

### BrdU pulse-chase experiment

1mg/ml BrdU (Sigma, #B5002), in combination with 1% sucrose, was administered to mice via their drinking water for 14 days. BrdU-containing drinking water was exchanged every three days. Mice were sacrificed after 14 days of continuous BrdU labelling to assess the initial labelling efficiency and after a chase period of 10 and 21 days to assess label retention. Intestines were removed, flushed with ice-cold PBS, fixed with 4% PFA for 3h at 4°C, cryoprotected by a sucrose gradient and embedded in OCT. Cryosection imaging was performed using a Leica SP5 confocal microscope and cells were counted manually.

### 5-FU treatment

Intestinal injury was induced by injecting two intraperitoneal doses of 5-fluorouracil (5-FU, 100 mg/kg, Sigma) over a 48h period. Mice were sacrificed 48h or 26 days after the last 5-FU dose and intestinal tissue was analyzed by immunohistochemistry and FACS. To assess the replication rate in the small intestine, 5-Ethynyl-2-deoxyuridine (EdU) (Thermo Fisher Scientific, A10044) was administered as an i.p. injection at 100 μg/g body weight from a 10 mg/ml stock in sterile PBS. Mice were sacrificed 4 h post EdU administration. Lgr5^hi^ and mTmG cells were isolated by flow cytometry and cytospun on glass slides. EdU staining was performed on cytopun cells using the Click-iT Staining Kit (Invitrogen, #C10340) according to the manufacturer’s instructions.

### Western blot analysis

For Western blot analysis, FVR^+^ and FVR^-^ cells were FAC-sorted and lysed in RIPA buffer (50mM Tris pH7.5, 150mM NaCl, 1mM EDTA, 1% Igepal, 0.1% SDS, 0.5% Sodiumdeoxycholate) containing phosphatase inhibitor (Sigma-Aldrich, P5726, P0044) and proteinase inhibitor (Sigma-Aldrich, P8340). Cell lysates were resolved by SDS-PAGE, transferred to PVDF membrane (Biorad) and incubated with the following primary antibodies: rabbit anti-pJnk (1:1000, NEB, #4668); rabbit anti-Jnk (1:1000, NEB, #9258); mouse anti-Gapdh (Merck Biosciences, CB1001); rabbit anti mTor (Cell Signaling, #2972); rabbit anti-pmTor (Ser2448) (Cell Signaling, #5536). Protein bands were visualized using horseradish peroxidase (HRP)-conjugated antibody, goat anti-mouse HRP (Dianova, 115-036-062) or goat anti-rabbit HRP (Dianova, 111-036-045) and chemiluminescence reagent (Millipore). The bands were quantified using ImageJ.

### Flow cytometry

For gene expression (microarray, single-cell RNAseq/qRT-PCR, qRT-PCR) analysis, Western Blot, and single-cell culture, crypt cells were sorted using FACS-Aria III (BD Bioscience) with a 100 μm nozzle. For all experiments, single cells were gated according to their FSC-A (front scatter area) and SSC-A (side scatter area). Singlets were gated dependent on the FSC-W (front scatter width) and FSC-H (front scatter height) and dead cells were excluded using the 7-AAD marker (eBioscience, #00-6993-50).

### Crypt isolation, crypt culture in matrigel, single-cell preparation for FACS, intestinal single-cell culture

Isolation and culture of small intestinal crypts and organoid culture was performed as previously described^40^. Briefly, intestines were harvested and washed with PBS. Villi were scraped away using coverslips. The remaining tissue was cut into 2cm pieces and incubated in 2mM EDTA/PBS for 35min at 4°C. Finally, crypts were collected by shaking. For Wnt-stimulation, isolated crypts were cultured in growth factor-free matrigel (BD Bioscience #356231) overlaid with medium containing 50ng/ml EGF (Life technologies PMG8043)/100ng/ml mNoggin (Peprotech, #250-38)/1μg/ml mR-spondin1 (R&D sytems, #2474-RS-050) (ENR) in the presence of 10μM Rock-inhibitor (Sigma, Y0503). Crypts were plated in 24-well plates at a density of 400 crypts/40μl matrigel. Two days after plating, the medium was changed for Wnt stimulation to ENR containing 400ng/ml Wnt ligand (Wnt5a, R&D systems #645-WN-010; Wnt11, R&D systems #6179-WN-010). After 2 days culture with Wnt ligands, crypts were intensively washed with ice-cold PBS and lysed in QIAzol (Qiagen, #79306) for RNA isolation (Qiagen, #79306) and cDNA synthesis (Invitrogen, SuperScript VILO cDNA synthesis kit, #11754).

For single-cell preparation, the crypt pellet was resuspended in 1-1,5ml TrypLE (Life technologies, #12605), incubated on ice for 5min, followed by 5min incubation at 37°C in a water bath. Then, 6ml of crypt complete medium containing 10% FCS and 10μg/ml DNase were added, and cells were incubated for 5min at 37°C in a water bath. The cells were gently resuspended by pipetting up and down 10 times, 10ml FACS buffer (2% FCS, 2mM EDTA in PBS) was added and the cells were centrifuged at 300xg, 5min, 4°C. Cells were washed twice with FACS buffer and finally the cell pellet was re-suspended in 1-2ml FACS buffer containing 10μM Rock-inhibitor (Sigma, Y0503), and cells were passed through the 40μm cell strainer caps of FACS tubes.

Single-cell culture to assess organoid formation efficiency, was performed as described previously^41^. 6,000 cells/25μl matrigel (BD Bioscience #356231) were seeded in a 24-well and overlaid with medium containing ENR, 10μM Rock-inhibitor (Sigma, Y0503), 1mM Valproic acid (Sigma, PHR1061) and 3μM CHIR99021 (Stemgent). Medium was changed every two days. VPA and CHIR99021 were added for the first 6 culture days. Bright-field images were acquired using a Zeiss microscope. Organoids were counted after 12 days culture.

### RNA isolation, qRT-PCR and microarray mRNA profiling

For gene profiling and qRT-PCR cells were directly sorted into Qiazol lysis reagent (Qiagen, #79306) and total RNA was extracted using the miRNeasy Micro kit (Qiagen, #217084), RNA integrity was checked using Agilent 2100 Bioanalyzer (Agilent RNA 6000 Pico Kit) and cDNA was amplified with the Ovation PicoSL WTA System V2 in combination with the Encore Biotin Module (Nugen, USA). Amplified cDNA was hybridized on Affymetrix Mouse Gene 1.0 ST arrays (FVR) or Affymetrix Mouse Gene 2.0 ST arrays (Lgr5^hi^/mTmG). Staining and scanning was performed according to the Affymetrix expression protocol, including minor modifications as suggested in the Encore Biotin protocol. For FVR data Expression Console (v.1.3.0.187, Affymetrix) was used for quality control and to obtain annotated normalized RMA gene-level data (standard settings including median polish and sketch-quantile normalisation). Lgr5^hi^/mTmG data were RMA normalized using R/Bioconductor package oligo (version 1.38.0) and probesets were annotated using the R/Bioconductor package mogene20sttranscriptcluster.db (version 8.5.0). Differential expression analyses were performed with the R environment for statistical computing (R Development Core Team, http://www.R-project.org/) by using the limma package (version 3.30.7) and P-values were adjusted for multiple testing by Benjamini-Hochberg correction. A gene was considered as differentially expressed if the adjusted p-value (FDR) was below a threshold of 0.05 (for FVR) or <0.01 (for Lgr5^hi^/mTmG; an additional filter for fold-change>2x was applied). Heatmaps were drawn using the pheatmap library for R. Expression is scaled row-wise and the colours range from dark blue (low expression) to orange (high expression). Functional enrichments were conducted using the GOstats package for R.

### TaqMan qRT-PCR

TaqMan qRT-PCR was performed under standard conditions using ViiA7 (Applied Biosystems) and TaqMan Fast Advanced Master Mix (Applied Biosystems, #4444557) or TaqMan Universal Master Mix II (Applied Biosystems, #4440040) for amplified cDNA. Samples were normalized to housekeeping genes: 18S ribosomal RNA (*RN18S*) and glyceraldehyde 3-phosphate dehydrogenase (*Gapdh*).

Taqman probes (Applied Biosystems): *Lyz1*, Mm00657323_m1; *Chga*, Mm00514341_m1, *Gapdh*, Mm99999915_g1; *RN18S*, Mm03928990_g1; *Mki67*, Mm01278617_m1; *Cdkn1a* (p21), Mm04205640_g1; *Muc2*, Mm01276696_m1; *Ccnd1*, Mm00432359_m1; *Ror2*, Mm01341765_m1/ Mm00443470_m1; *Fltp*, Mm01290543_g1; *Fltp*, Mm01290541_m1 *Prickle1*, Mm01297035_m1; *Dvl2*, Mm00432899_m1; *Celsr1*, Mm00464808_m1; *Fzd6*, Mm00433387_m1; *Jun*, Mm00495062_s1.

### Single-cell gene expression analysis by microfluidic qRT-PCR

Small intestinal crypt cells were kept cold and sorted using FACS-Aria III (BD Bioscience). Doublets were excluded and dead cells were excluded using 7-AAD (eBioscience, #00-6993-50). The pre-amplification solution in 96-wells included 5μl of a master mix containing 1.2μl 5x VILO reaction mix (Invitrogen, #11754-050), 0.3μl 20U/μl SUPERase-In (Ambion, #AM2694), 0.25μl 10% NP40 (Thermo Scientific, #28324), 0.25μl RNA spikes mix (Fluidigm, #100-5582) and 3 μl nuclease-free water (Promega, #P119C). Cells were lysed by incubation at 65°C for 90s and RNA was transcribed into cDNA by adding 1μl of RT mix solution containing 0.15μl 10x SuperScript enzyme mix (Invitrogen, #11754-050), 0.12μl T4 Gene 32 Protein (New England BioLabs, #M0300S) and 0.73 μl nuclease-free water and RT cycling (25°C for 5min, 50°C for 30min, 55°C for 25min, 60°C for 5min and 70°C for 10min). Target specific cDNA amplification was performed by adding 9μl reaction mix containing 7.5μl TaqMan PreAmp Master Mix (Applied Biosystems, #4391128), 0.075μl 0.5M EDTA, pH 8.0 (Invitrogen, #Am9260G), 1.5μl 10x outer primer mix (500nM) (see Supplementary Table 6) and 20 cycles of denaturation for 5s at 96°C and 4 min annealing/extension at 60°C following an enzyme activation step at 95°C for 10min. Exonuclease I treatment was performed to clean up the reaction by adding a 6μl reaction mix containing 0.6μl reaction buffer, 1.2μl Exonuclease I (New England BioLabs, #M0293S) and 4.2μl nuclease-free water.

Amplified single-cell cDNAs were analysed with gene specific inner primer pairs (Supplementary Table 6) and SsoFast EvaGreen Supermix with Low ROX (Bio-Rad Laboratories, #172-5210) using the 96 × 96 Dynamic Array on the BioMark System (Fluidigm). Ct values for each gene in each cell was calculated using BioMark <u>Real-Time PCR</u> Analysis software (Fluidigm).

### Computational analyses of single-cell qRT-PCR data

Subsequent data processing and analysis of single cell qRT-PCR data was performed in R (http://www.r-project.org/). We manually removed 8 cells due to technical problems (pipetting error) during experimental processing. Samples were then corrected for deviating sample dilution of PCR runs. We followed the normalization procedure as suggested in9. Briefly, we subtracted Ct values from the assumed limit of detection of the BioMark (LOD=30). As quality control measure and reference we used the expression values of the three most robustly expressed housekeeping genes Rn18S, ActB and Hsp90. We excluded all cells that did not express all three housekeepers as well as cells for which the mean of the three housekeepers was ±3 s.d. from the mean of all cells. ΔCt values were then normalized on a cell-wise basis to the mean expression of the three housekeeping genes. The minimum of the normalized data - 2 was then assigned as a ΔCt value where a gene was not detected (failed qPCR runs). 595 of 672 sorted cells were retained for further analysis. Foxa1, Foxa2, GFP did not amplify correctly in one run so these genes have been excluded from the analysis as were all housekeeping genes (Rn18S, ActB, Hsp90, Uba52, Gapdh, Rpl37). We quantified three synthetic RNAs of different concentrations to explore its use as a reference. The signals of the RNA spikes were too strong and not quantifiable in all experimental runs and were therefore removed before further analyses. *Fltp* was detected in the negative control of Neurog3 cells and therefore had to be excluded from multivariate analyses. For the other cell types *Fltp* measurements were correct. In total, we used ΔCt values of 80 genes.

### Single-cell qPCR analysis

All further analysis of single-cell qPCR data was performed using Scanpy (v.1.0.4). The singlecell neighborhood graph was computed on the 15 first principal components with a local neighborhood size of 5 (pp.pca and pp.neighbors) and UMAP was run for visualization (tl.umap). Louvain-based clustering at a resolution of 0.8 was used for subtype identification which were annotated based on the expression of known marker genes (pp.louvain). Genes characteristic for each subtype were identified using a wilcoxon-ranksum test (tl.rank_gene_groups). Top 5 ranked genes were considered for plotting. For the reconstruction of lineage relationships and differentiation trajectories we used PAGA (tl.paga) and diffusion pseudotime (dpt, tl.dpt). We first applied PAGA to find the branching into the EEC and Paneth cell lineage. For each lineage (branch in PAGA) we then arranged cells by their pseudotemporal order inferred from dpt (pl.paga_path). The root is represented by a cell in the ISC population, defined as Lgr5^+^ cells expressing the stem cell markers (*Lgr5, Olfm4, Ascl2, Axin2* and *Prom1*) but without specific lineage markers (*Lyz1, Mmp7, Atoh1, Dll4, Dll1* or *Sis*). Random variation of the root within the stem cell population did not substantially change dpt. Expression values along a trajectory are plotted as the smoothed average over n cells using a sliding window with Gaussian noise as implemented in pl.paga_path (n as indicated in figure legend).

### Single-cell RNA sequencing: RNA preparation, library generation and sequencing

Samples from SI crypts were prepared as described above under section crypt isolation and flow cytometry (Supplementary Table 2). For rare lineage enrichment live crypt cells were mixed with reporter positive cells at different ratios. Number of dead cells was estimated by trypan blue staining and sorted cells were counted. Single-cell libraries were generated using the Chromium™ Single cell 3’ library and gel bead kit v2 (10X Genomics, #120237) according to the manufacturer’s instructions. Briefly, cells were loaded onto a channel of the 10X chip to generate Gel Bead-in-Emulsions (GEMs). These underwent reverse transcription to barcode RNA before cleanup and cDNA amplification. Afterwards, enzymatic fragmentation and attachment of 5’ adaptor and sample index was performed. Libraries were sequenced on the HiSeq4000 (Illumina) with 150 bp paired-end sequencing of read 2.

### Computational analysis

#### Pre-processing of droplet-based scRNAseq data

De-multiplexing and alignment to mm10 mouse genome, identification of unique molecular identifiers (UMI) and barcode filtering was performed using the ‘CellRanger’ toolkit (version 2.0.0) provided by 10X Genomics. We performed a further barcode (=cell) selection step and additionally included cells with more than 1000 expressed genes, where a gene is counted as expressed if we found at least one UMI mapped to it. We further filtered cells with a fraction of counts from mitochondrial genes > 10% indicative for stressed or dying cells.

We removed doublets by computing the doublet score from *scrublet*^42^ on the UMI count matrix separately for every sample. Using a threshold of 0.4, we removed 1651 cells from the analysis.

Cells from all samples were log-normalised and batch corrected using ComBat as Python implementation. Please note that we observed an overrepresentation of Paneth cells in the FVR-enriched samples (with 50% and 90% enrichment, resp.), where we subsampled the Paneth cell populations to 15% to fit the cell-type distribution of all other samples before we corrected with ComBat. Then, we used the pre-computed regression coefficients to correct for batch effects in the filtered cells and merged them with the full data set. Further, we fixed all zero values to remain zero in order to preserve the support of the count data. We computed the top 2000 highly variable genes based on mean and dispersion (pp.filter_genes_dispersion in SCANPY v. 1.3.1 in Python 3.6 with the flavor ‘cell_ranger’ to compute normalised dispersions^43^).

Further, we corrected for the library size by scaling the reads per cell to the factor 100,000 (pp.normalize_per_cell in SCANPY v. 1.3.1 in Python 3.6).

#### Dimension reduction

We performed our analyses with SCANPY^44^ v. 1.3.1 in Python 3.6. We used a UMAP^45^ to represent the data in the two-dimensional embedding and data visualisation (tl.umap). This was created based on a PCA-space with n=50 components, and the k-nearest neighbour graph on the PCA-space with k=30 (tl.pca and pp.neighbors). PCA etc. was computed on the scaled and normalised data with 2,000 highly variable genes.

#### Clustering and cell type annotation

We determined the clustering and cell type annotation for the control samples as follows. We inspected first marker gene expression for respective major cell types and computed gene scores using known marker genes (tl.score_genes, based on ref.^46^). Analogously, we computed a cell cycle score to determine the respective cell-cycle phase state of the cells (tl.score_genes_cell_cycle, based on ref.^46^). Then, we performed clustering using the louvain algorithm^47^ (tl.louvain with default resolution parameter 1.0) and found 18 clusters. Here, we annotated and merged clusters again according to the gene scores and marker gene expression. Here, we also identified 1589 immune cells, which were distinct from the remaining cells. Subsequently, we inspected all main clusters for substructure and resolved it further based on marker gene expression. Here, we used louvain clustering with resolution parameters ~1-2 and merged the subclusters again according to marker gene expression (hierarchical ‘split-and-merge’ approach). Finally, we annotated 7 major cell types (ISC, Enterocytes, Goblet cells, Paneth cells, Enteroendocrine cells and Tuft cells), subdivided them into progenitor and mature cells. In the ISC population, we identified a PC-primed ISC population that was more similar to the Paneth progenitor population.

For the mutant samples, we employed a k-nearest neighbour approach to match every cell to the corresponding cluster (k=30), i.e. we derived the cell identity of the mutants from the cell identity of neighbouring cells of the control samples, which we annotated beforehand. Again, we identified 2289 immune cells, which we removed from analysis.

#### Differential expression analysis

We used the limma package^48^ (version 3.34.9 in R 3.4.3) to study differential expression. First, we excluded the FVF enriched samples and mutant samples. In order to determine differentially expressed genes between ISCs and progenitor populations, we tested pairwise progenitor populations vs ISC (without Paneth primed ISCs) and Paneth progenitors vs Goblet progenitors. Please note that differential expression was performed on the log-normalised (not batch-corrected) data, where we included the sample information (i.e. batch) as covariate. In addition, we excluded all genes with mean expression < 0.05. We considered only significant genes (FDR<0.05) with logFC>0.05. For the analysis of transcription factors, we filtered differentially expressed transcription factors (gene ontology ID GO:0003700 (transcription factor activity)) with the biomartr package^49^ (version 0.7.0 in R 3.4.3) with a p-value threshold p_adj_<10^-5^. Analogously, in order to determine differentially expressed genes between mutants and control samples (without FVF enriched samples), we tested every cluster separately with limma. Please note that differential expression was performed on the log-normalised (not batch-corrected) data, where we included the number of expressed genes as covariate, but not the sample information (i.e. batch) due to confounding of genetic condition and sample covariates. In addition, we excluded all genes with mean expression < 0.05. We considered only significant genes (FDR<0.05) with logFC>0.05. For the analysis of transcription factors, we filtered differentially expressed transcription factors (gene ontology ID GO:0003700 (transcription factor activity)) with the biomartr package (version 0.7.0 in R 3.4.3) and with an adjusted p-value threshold p_adj_<10^-5^ (Benjamini-Hochberg correction).

#### Identifying cell differentiation trajectories via graph abstraction

To derive cell trajectories, we computed a pseudotemporal ordering using diffusion pseudotime (DPT, tl.dpt in SCANPY)^50^. As the topology of the data is complex, we used partition-based graph abstraction (PAGA)^33^ to quantify the connections between the clusters (*i.e*. connections of ISCs to respective progenitors and mature cell types, tl.paga in SCANPY). We display all connections with a scaled connectivity of at least 0.05 (‘threshold’ parameter in pl.paga in SCANPY).

#### Gene Set Enrichment Analysis

We performed a gene set enrichment analysis based on both GO^51,52^ terms and KEGG^53^ terms using g:profiler^54^. In particular, we adapted the Python wrapper from V. Svensson (https://github.com/vals/python-gprofiler). We set the background to all expressed genes and removed all resulting gene sets with significance level p>=0.05. Further, we split the input data set by the sign of the log-fold change, such that we considered up-regulated and down-regulated gene sets separately in each set of differentially expressed genes. For visualisation of gene set significance, we abridged p-values at 10^-10^.

### Code and data accession

Custom R scripts of the single cell qRT-PCR bioinformatics analyses are available in a jupyter notebook upon request.

Microarray data have been submitted to NCBI/GEO (GSE94092). scRNAseq data have been submitted to NCBI/GEO (code upon request).

### Statistical analysis

No statistical methods were used to predetermine sample size. The experiments were not randomized and the investigators were not blinded to allocation during experiments and outcome assessment.

Statistical analysis was performed using GraphPad Prism 6 Software (GraphPad Software, USA). Data are expressed as values and were compared using unpaired t-tests or ANOVA, unless indicated otherwise.

---

**Supplementary Table 1**

Shows the differentially expressed genes in the FVR populations.

**Supplementary Table 2**

Overview of samples used for scRNAseq analysis.

**Supplementary Table 3**

Shows the enrichment of rare lineages with Fltp^ZV^ and FVF reporter mice.

**Supplementary Table 4**

Shows the genes used to determine the gene scores.

**Supplementary Table 5**

Genes used in single-cell qRT-PCR analysis.

**Supplementary Table 6**

Primer for single-cell qRT-PCR.

## Acknowledgements

We thank Kerstin Diemer, Jürgen Schultheiss, Ines Kunze, Anita Ludwig, and Anke Bettenbrock for excellent technical assistance, and Aurelia Raducanu and Pallavi Mahaddalkar for assistance with the FACS, and Henner Farin for teaching intestinal single cell culture. We are grateful to Jennifer Murdoch for providing *Celsr1^crsh^* animals and Helena Edlund for providing the Ngn3 antibody. We thank Ralph Böttcher, Matthias Tschoep and Steve Woods for critical reading the manuscript. This work was supported by an Emmy-Noether Fellowship and the European Union with the ERC starting grant Ciliary Disease and HumEn project. This work was funded by the Helmholtz Alliance ICEMED – Imaging and Curing Environmental Metabolic Diseases (H.L., J.B.) and through the Initiative and Networking Fund of the Helmholtz Association (H.L.). For financial support we would also like to thank the Helmholtz Society, Helmholtz Portfolio Theme ‘Metabolic Dysfunction and Common Disease’ (H.L. and J.B.), the Helmholtz Alliance ‘Aging and Metabolic Programming, AMPro’ (H.L., J.B.). Further, this projected was funded by ExNet-0041-Phase2-3 (“SyNergy-HMGU”) through the Initiative and354Network Fund of the Helmholtz Association (H.L., F.J.T.), the German Research Foundation and the German Center for Diabetes Research (DZD e.V.) (H.L.).

## Authors Contributions

A.B. designed experiments, performed experiments, analysed data and wrote the manuscript. M.B. analysed single-cell qRT-PCR and RNAseq data. S.T. analysed single-cell qPCR data. M.S. analysed microarray data. A.A. performed western blot analysis of sorted cell populations. I.B. generated Ngn3-VF mouse line. S.S. analysed microarray data. M.I. and J.B. performed the microarrays and data analysis. C.Z. and W.E. contributed to single cell qPCR experiment and discussions. A.C.S., F.M.V. and O.E. provided single-cell qRT-PCR resources and contributed to discussions. F.J.T. supervised M.B., S.T. and S.S. and analysed single-cell RNAseq and qRT-PCR data. H.L. helped in study design, analysis and writing and acquired financial support.

---

**Extended Data Figure 1.**
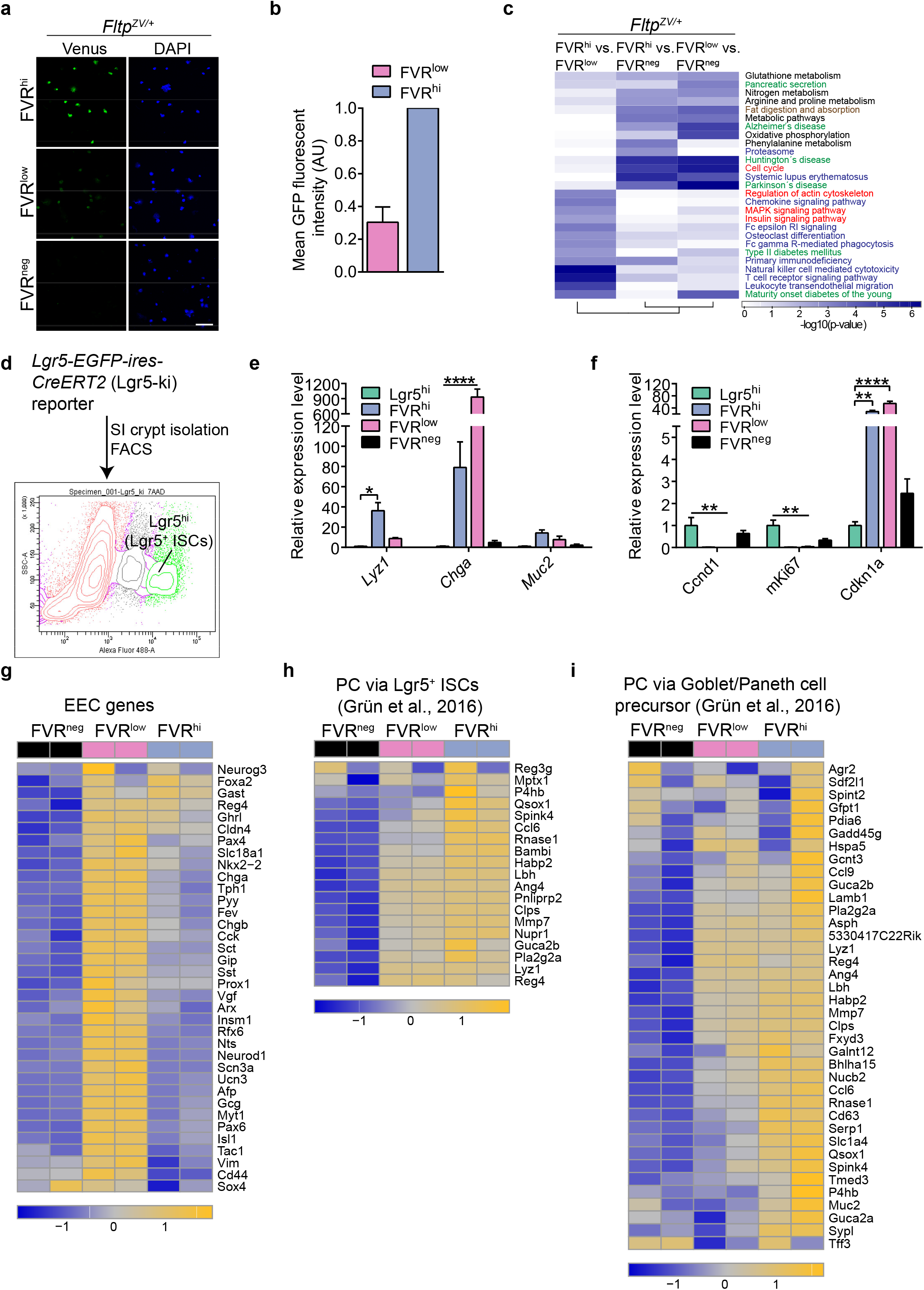
FVR labels all Paneth and enteroendocrine subtypes. **a,** LSM images of immune-stained flow-sorted FVR^hi/low/neg^ SI crypt cells isolated from *Fltp^ZV/+^* mice showing that three distinct crypt cell populations are distinguishable by FVR activity. FVR (Venus, green), DAPI (blue, nuclei). Scale bar, 75 μm. **b,** Relative mean fluorescent intensity of GFP (Venus) in FVR^low^ and FVR^hi^ cells isolated from *Fltp^ZV+^* cells by flow cytometry. *n* (independent experiments) = 6. Error bar represent SD. **c,** Heatmap depicting the p-values determined by Fisher’s exact test of Kyoto Encyclopedia of Genes and Genomes (KEGG) pathways which were significantly enriched in at least one of the populations. Pathways associated with neuroendocrine (green), immuno-regulatory function (blue), actin cytoskeletal regulation and signalling (MAPK/ Insulin, red) as well as cellular metabolism (black) and the cell cycle (red) are indicated. **d,** FACS plot depicting the experimental strategy to obtain Lgr5^hi^ cells (intestinal stem cells, ISCs) from *Lgr5-EGFP-ires-CreERT2* (Lgr5-ki) mice. **e, f,** Confirmation of the microarray results. Comparison of the crypt cell populations distinguishable by FVR activity with Lgr5^hi^ ISCs by qRT-PCR, for the expression of intestinal lineage markers (*Lyz1*, Paneth cells; *Chga*, enteroendocrine cells; *Muc2*, goblet cells) (**e**) and proliferation markers (*Ccnd1*, Ki67) and cell-cycle exit marker (*Cdkn1a*) (**f**). *n* (independent experiments) = 3. Error bars represent SEM. Two-tailed Student’s *t*-test, \**P*<0.05, \*\**P*<0.01, \*\*\*\**P*<0.0001. **g, h, i,** Heatmaps depicting the enrichment of the enteroendocrine cell (EEC) gene signature in the FVR^low^ population (**g**), and the enrichment of genes expressed in Paneth cells (PC) formed via Lgr5^hi^ ISCs (**h**) or via a Goblet/Paneth cell precursor (**i**) in FVR^hi^ cells. Gene signatures are derived from Grün et al., 2016. FVR labels all reported EEC subsets^29^. Expression is scaled row-wise and the colours range from dark blue (low expression) to orange (high expression) and represent normalized expression (row z-score).

**Extended Data Figure 2.**
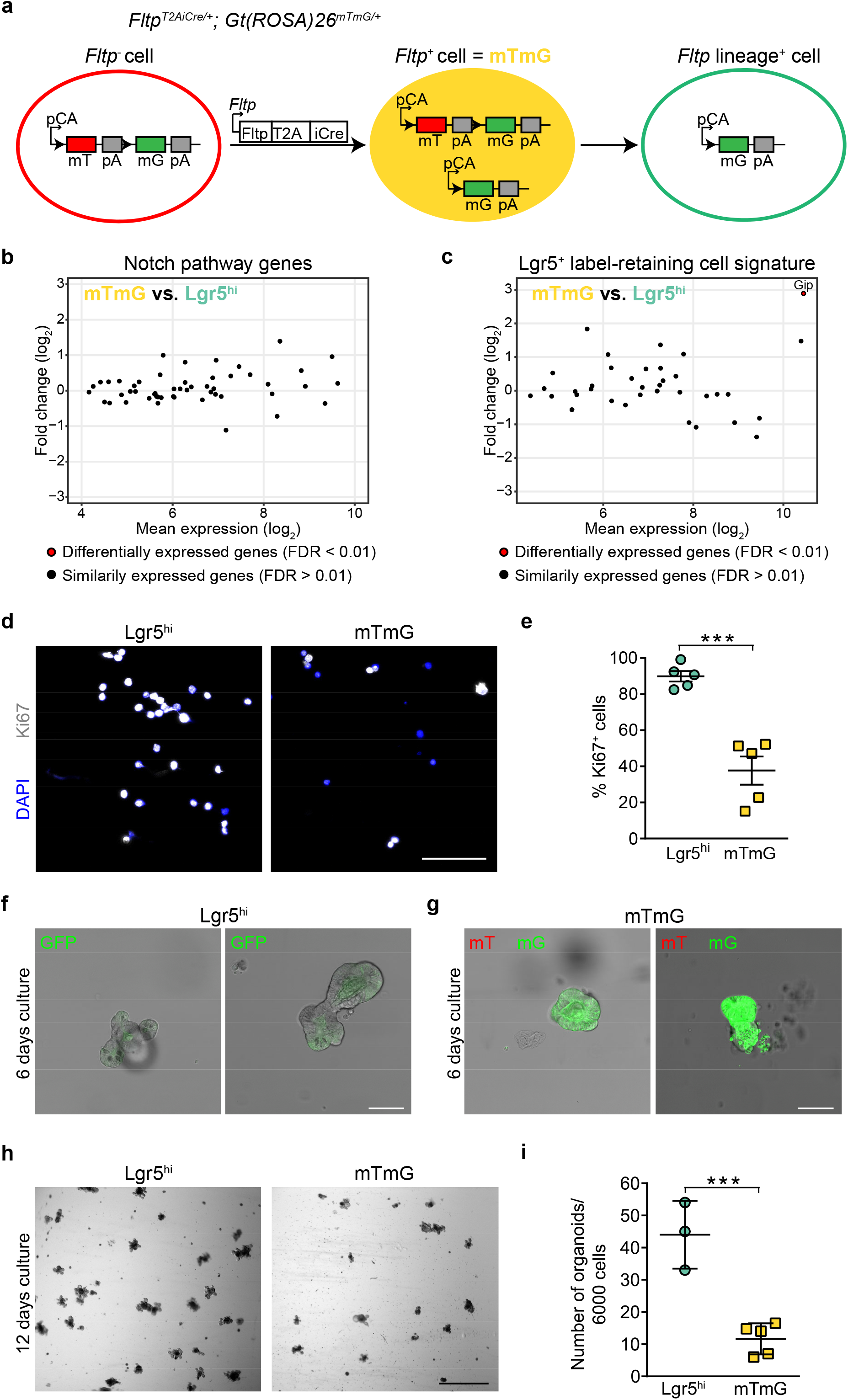
*Fltp^+^* cells possess limited organoid forming capacity *in vitro*. **a,** Schematic depicting the *Fltp^T2AiCre/+^; Gt(ROSA)26^mTmG/+^* lineage-tracing model. Fltp lineage^-^ cells (mT, red) convert into Fltp lineage^+^ cells (mG, green) upon Fltp-promoter driven Cre expression via an intermediate (mTmG, yellow) cell state. mTmG cells represent cells that started to express *Fltp*. **b,** MA-plot comparing the expression of Notch pathway genes (KEGG mmu 04330) in mTmG and Lgr5^hi^ (ISCs) cells. The y-axis depicts the fold change in log2 and the x-axis depicts the mean log2 expression value. No gene is significantly regulated. *n* (microarray samples) = 6 for Lgr5^hi^ ISC. *n* (microarray samples) = 4 for mTmG. **c,** MA-plot comparing the expression of Lgr5^+^ label-retaining cell signature genes^7^ in mTmG and Lgr5^hi^ (ISCs) cells. Differentially expressed genes (FDR < 0.01) are shown in red. The y-axis depicts the fold change in log2 and the x-axis depicts the mean log2 expression value. *Gip* is the only significantly regulated gene. *n* (microarray samples) = 6 for Lgr5^hi^ cells. *n* (microarray samples) = 4 for mTmG. **d, e,** Representative LSM images showing Lgr5^hi^ and mTmG cells stained for Ki67 (white) and DAPI (blue, nucleus) (**d**) and quantification of Ki67 positive cells (**e**). *n* (mice) = 5 for Lgr5^hi^ cells. *n* (mice) = 5 for mTmG cells. Error bars represent SEM. Two-tailed Student’s *t*-test, \*\*\**P*<0.001. Scale bar, 75μm. **f, g,** Representative LSM images showing live organoids derived from single Lgr5^hi^ ISCs (**f**) and mTmG cells (**g**). Scale bars, 75μm. **h, i,** Representative bright-field images of 6,000 FACS-purified mTmG and Lgr5^hi^ cells cultured for 12 days (**h**) and quantification of organoid number (**i**). *n* (independent experiments) = 3 for Lgr5^hi^ cells. *n* (independent experiments) = 5 for mTmG cells. Error bars represent SD. Two-tailed Student’s *t*-test, \*\*\**P*<0.001. Scale bar, 5 mm.

**Extended Data Figure 3.**
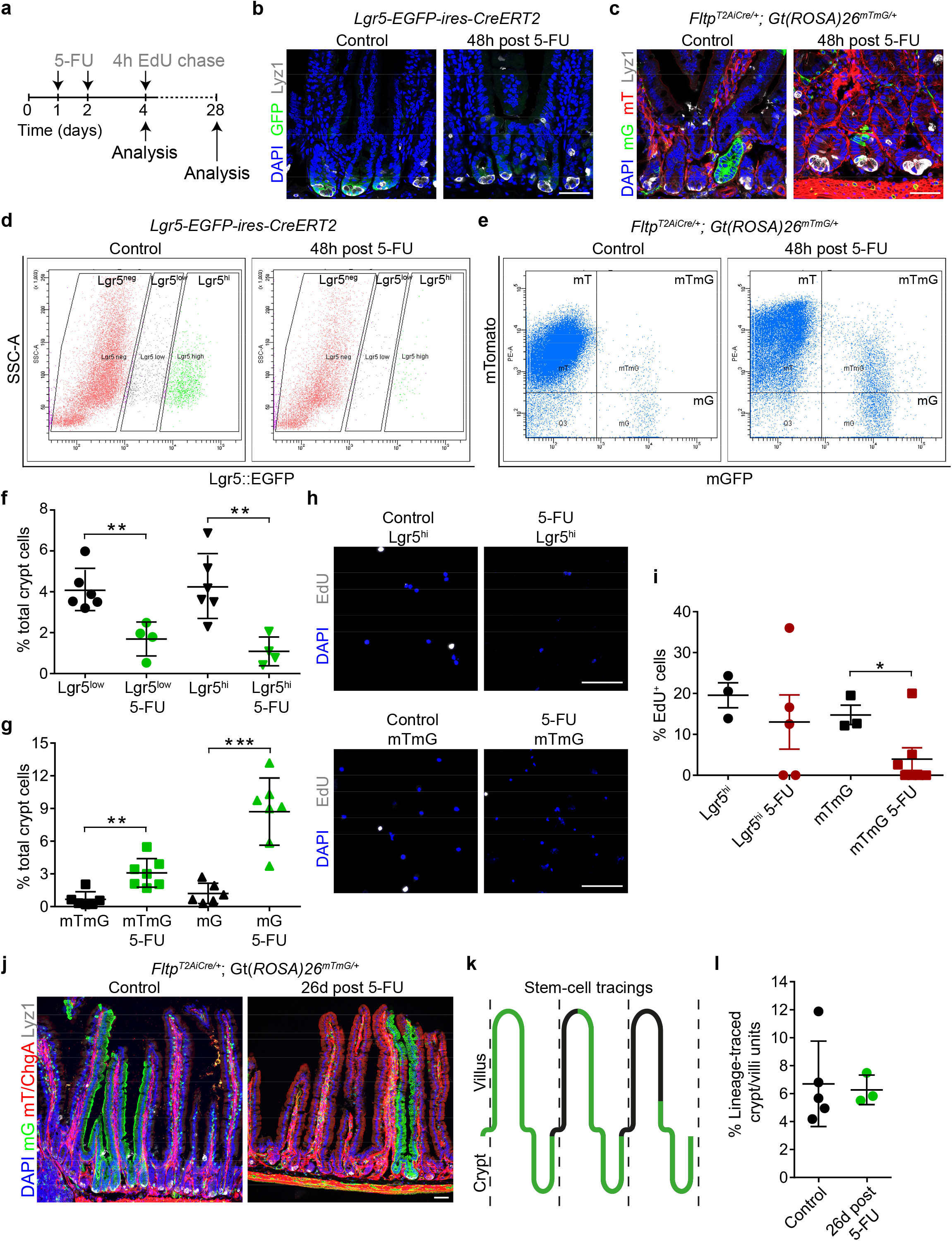
*Fltp^+^* cells possess limited multi-lineage potential *in vivo*, are resistant to chemical injury but do not contribute to regeneration after intestinal injury. **a,** Experimental setup to assess stress response of *Fltp^+^* cells (mTmG) compared to Lgr5^hi^ ISCs. Mice were treated with two consecutive doses of 5-FU (100 mg/kg/day). Lgr5^+^ cells and Fltp^+^ mTmG cells were analysed by flow cytometry 48 h and 24 days after the final 5-FU dose. Proliferative activity was determined after a 4 h chase of 5-ethynyl-2’-deoxyuridine (EdU) 48h after the final 5-FU dose. **b, c,** Representative LSM images of duodenal sections from Lgr5-ki (**b**) and *Fltp^T2AiCre/+^; Gt(ROSA)26^mTmG/+^* mice (**c**) 48h post 5-FU treatment. Sections were stained for DAPI (blue, nucleus), GFP (Lgr5^+^ cells, green) and Lyz1 (white, Paneth cells) (**b**) or mT (Fltp lineage^-^, RFP, red), mG (Fltp lineage^+^, GFP, green), and Lyz1 (white, Paneth cells) (**c**). Scale bars, 75μm. **d-g,** Representative FACS plots of Lgr5-GFP cell populations (**d**) and mT/mTmG/mG cell populations (**e**) from untreated and 5-FU treated mice and quantification of frequencies referred to total crypt cells (**f, g**) 48h post 5-FU treatment. *n* (mice) = 6 for untreated Lgr5-ki mice. *n* (mice) = 4 for treated Lgr5-ki mice. *n* (mice) = 6 for untreated *Fltp^T2AiCre/+^; Gt(ROSA)26^mTmG/+^* mice. *n* (mice) = 7 for treated *Fltp^T2AiCre/+^; Gt(ROSA)26^mTmG/+^* mice. Error bars represent SEM. Two-tailed Student’s *t*-test, \*\**P*<0.01, \*\*\**P*<0.001. **h, i,** Representative LSM images (**h**) and quantification (**i**) of 5-ethynyl-2’-deoxyuridine (EdU) positive (white) Lgr5^hi^ and mTmG cells after a 4 h chase of 5-ethynyl-2’-deoxyuridine (EdU) 48h after the final 5-FU dose. DAPI (blue) stains the nucleus. *n* (mice) = 3 for untreated Lgr5-ki mice. *n* (mice) = 5 for treated Lgr5-ki mice. *n* (mice) = 3 for *Fltp^T2AiCre/+^; Gt(ROSA)26^mTmG/+^* mice. *n* (mice) = 7 for treated *Fltp^T2AiCre/+^; Gt(ROSA)26^mTmG/+^* mice. Error bars represent SEM. Two-tailed Student’s *t*-test, \**P*<0.1. Scale bars, 75μm. **j, k, l,** Representative LSM images showing immune-stained duodenal sections from untreated and 5-FU treated *Fltp^T2AiCre/+^; Gt(ROSA)26^mTmG/+^* mice. DAPI (blue) stains nuclei, mGFP (green) labels Fltp lineage^+^ cells, mTomato (red) marks Fltp^-^ cells, Lyz1 (Paneth cells, white) and ChgA (enteroendocrine cells, red) (**j**). Scheme showing the mGFP expression pattern in crypt-villi units that are considered as stem-cell tracings (mGFP expression in all intestinal lineages in crypt-villi units) (**k**) and quantification of stem-cell tracings from sections (**l**). *n* (mice) = 5 for untreated mice. *n* (mice) = 3 for 5-FU treated mice. Scale bar, 75 μm. Error bars represent SD.

**Extended Data Figure 4.**
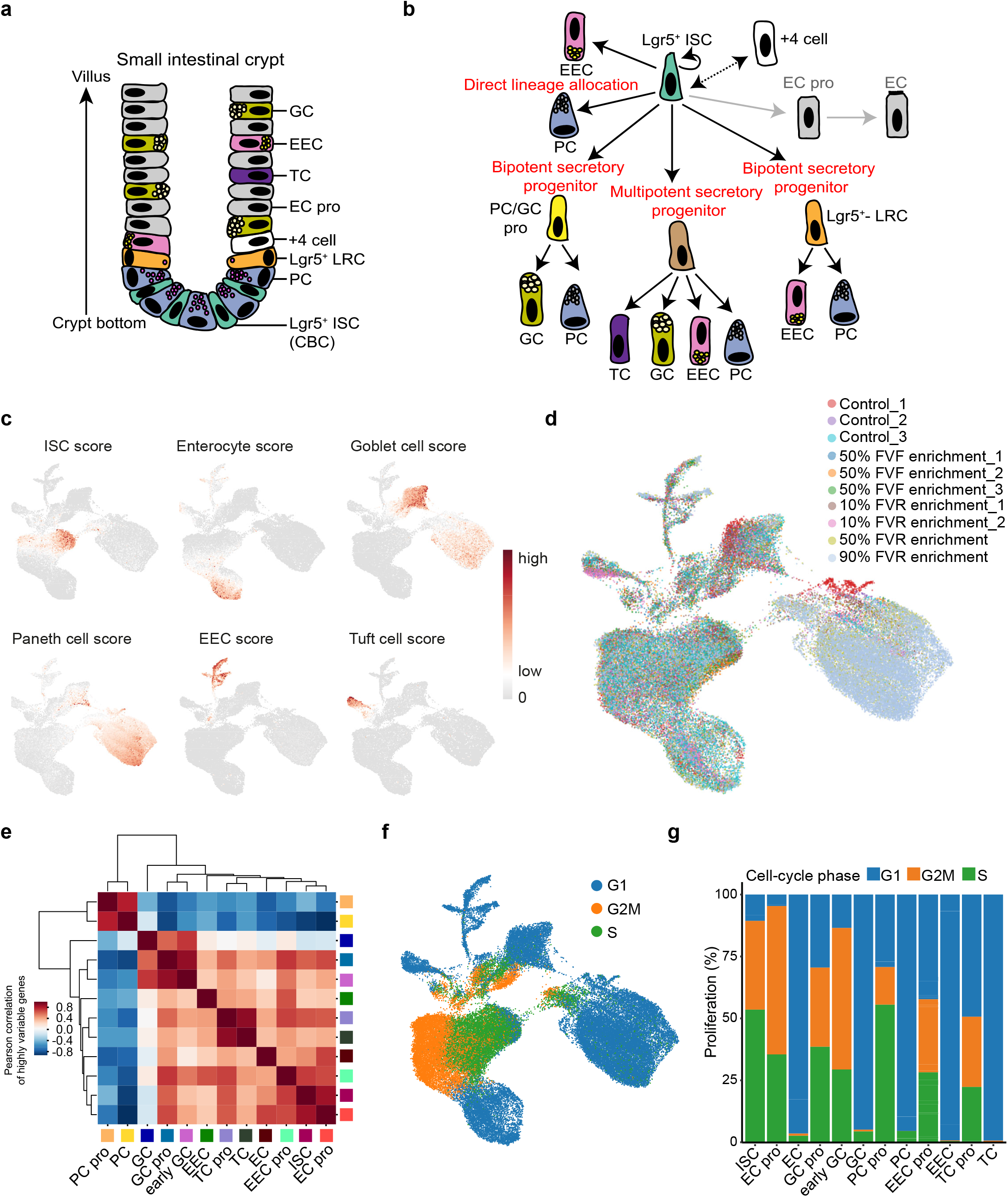
Identification of progenitors for each intestinal lineage by scRNAseq. **a,** Scheme of a SI crypt depicting all described intestinal epithelial cell types. **b,** Overview of current models of secretory and absorptive fate specification in the gut. Wnt/β-catenin dependent Lgr5^+^ ISCs self-renew or give rise to absorptive or multi-/ bipotent secretory progenitors that further differentiate into enterocytes or Paneth cells (PCs), enteroendocrine cells (EECs), goblet cells, tuft cells, respectively. A direct differentiation route from ISCs has been described for subtypes of Paneth and enteroendocrine cells. +4 reserve stem cells might correspond to Lgr5^+^ LRCs. **c,** UMAP depicting the gene score of major cell types (Supplementary Table 4). **d,** UMAP depicting all samples from control wildtype (no enrichment) and reporter mice (rare lineage enrichment) used in this study. **e,** Correlation map showing highly variable genes between progenitor cell types. **f,** UMAP depicting the cell-cycle score for each cell. **g,** Bar plot showing proliferation score of the progenitor and mature cell types.

**Extended Data Figure 5.**
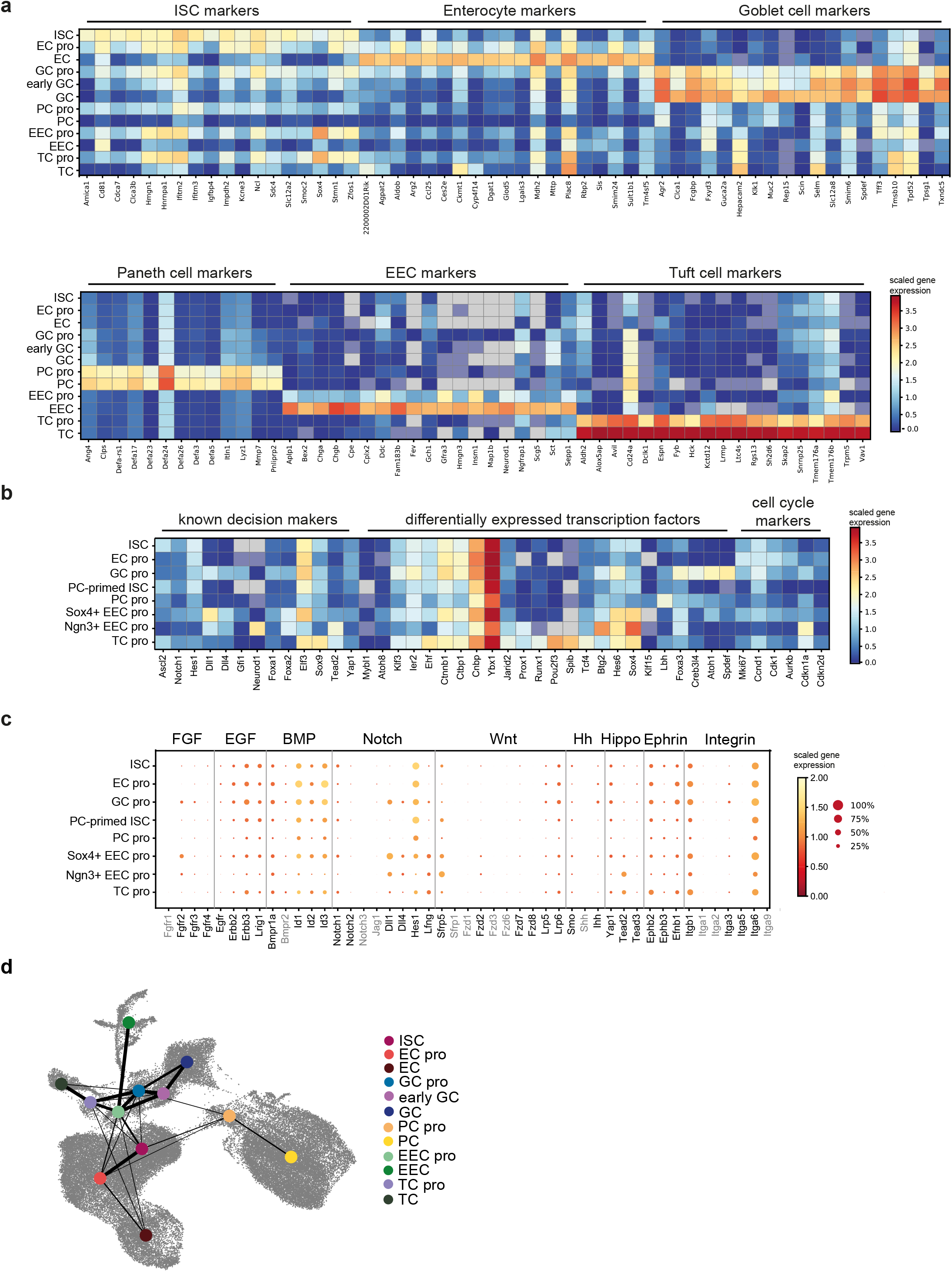
Intestinal progenitors are characterized by distinct gene expression patterns. **a,** Heatmap depicting the mean expression of marker genes in progenitor and mature cell populations. Distinct marker genes are already expressed in progenitor cells. **b,** Heatmap showing differentially expressed transcription factors between progenitors. **c,** Heatmap depicting signalling pathways involved in stem-cell maintenance, differentiation and cell positioning. Genes in black show significant expression differences. **d,** Abstracted graph of the PAGA model plotted on UMAP, cell-type cluster are reduced to colored dots. Graph abstraction reveals connections between cell types and are depicted as lines. Line thickness describes confidence level of the connection between two cell types (high confidence corresponds to high confidence). Connections below the threshold confidence of 0.05 are not displayed. All progenitor populations connect to ISCs.

**Extended Data Figure 6.**
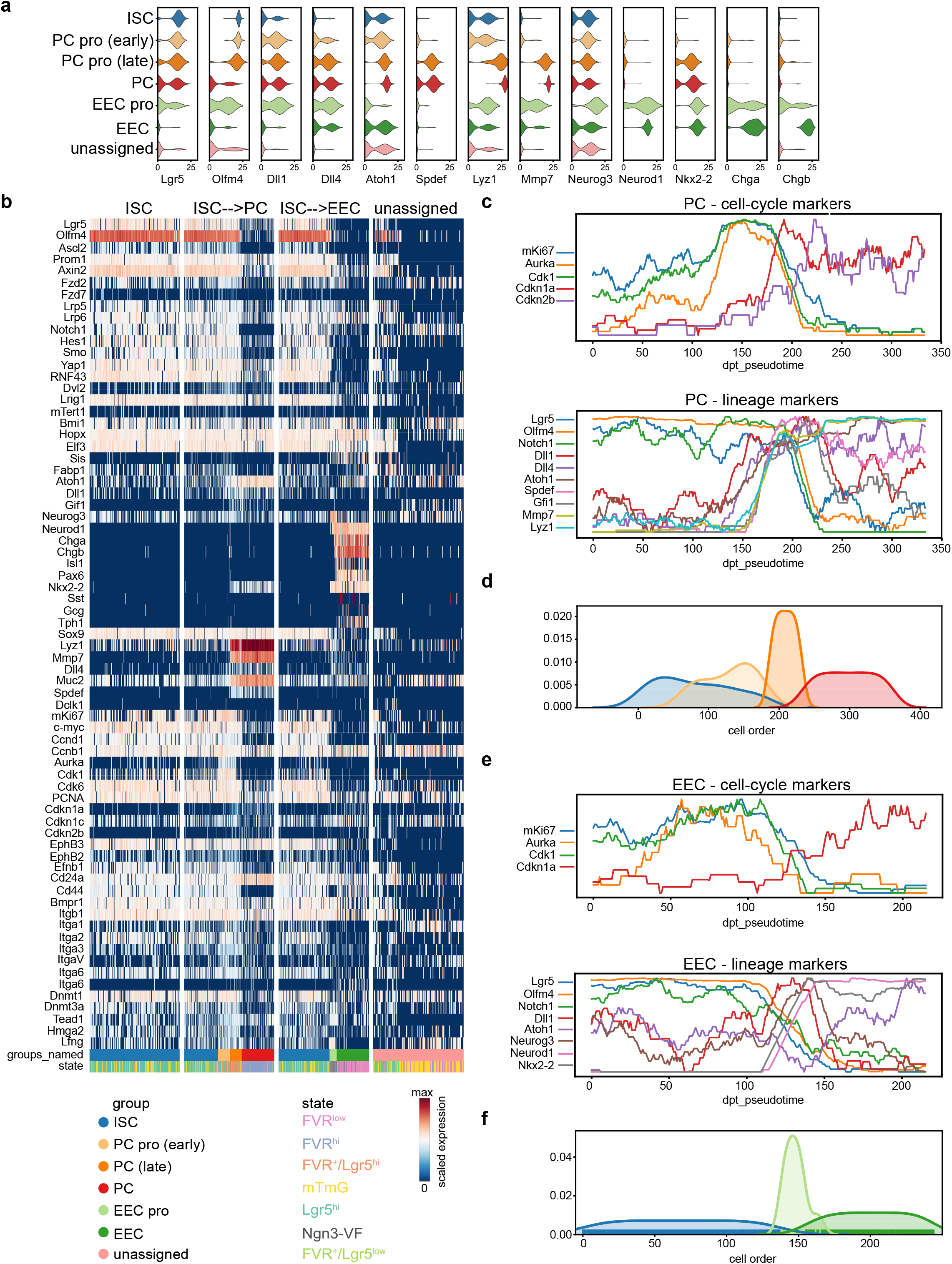
Pseudotemporal ordering of cells identifies unipotent transition states for EEC and PC lineage characterized by downregulation of stem-cell genes and co-expression of stem-cell and secretory lineage genes, respectively. **a,** Violin plots showing the distribution of selected marker genes from scqRT-PCR analysis in ISCs, and PC and EEC lineages. **b,** Heatmap of gene expression along the lineage-trajectories inferred by PAGA (separated by white vertical bars). Within each trajectory cells are ordered by DPT. Colour bars at the bottom indicate cell-type clusters and FACS states. **c,** Expression of cell-cycle (top) and lineage (bottom) marker genes along the PC lineage trajectory. Cells are ordered by DPT. Expression is shown as the running average over 30 cells. **d,** Cell density of cell-type clusters along the PC trajectory. Cells are ordered by DPT. **e,** Expression of cell-cycle (top) and lineage (bottom) marker genes along the EEC lineage trajectory. Cells are ordered by DPT. Expression is shown as the running average over 30 cells. **f,** Cell density of cell type clusters along the EEC trajectory. Cells are ordered by DPT.

**Extended Data Figure 7.**
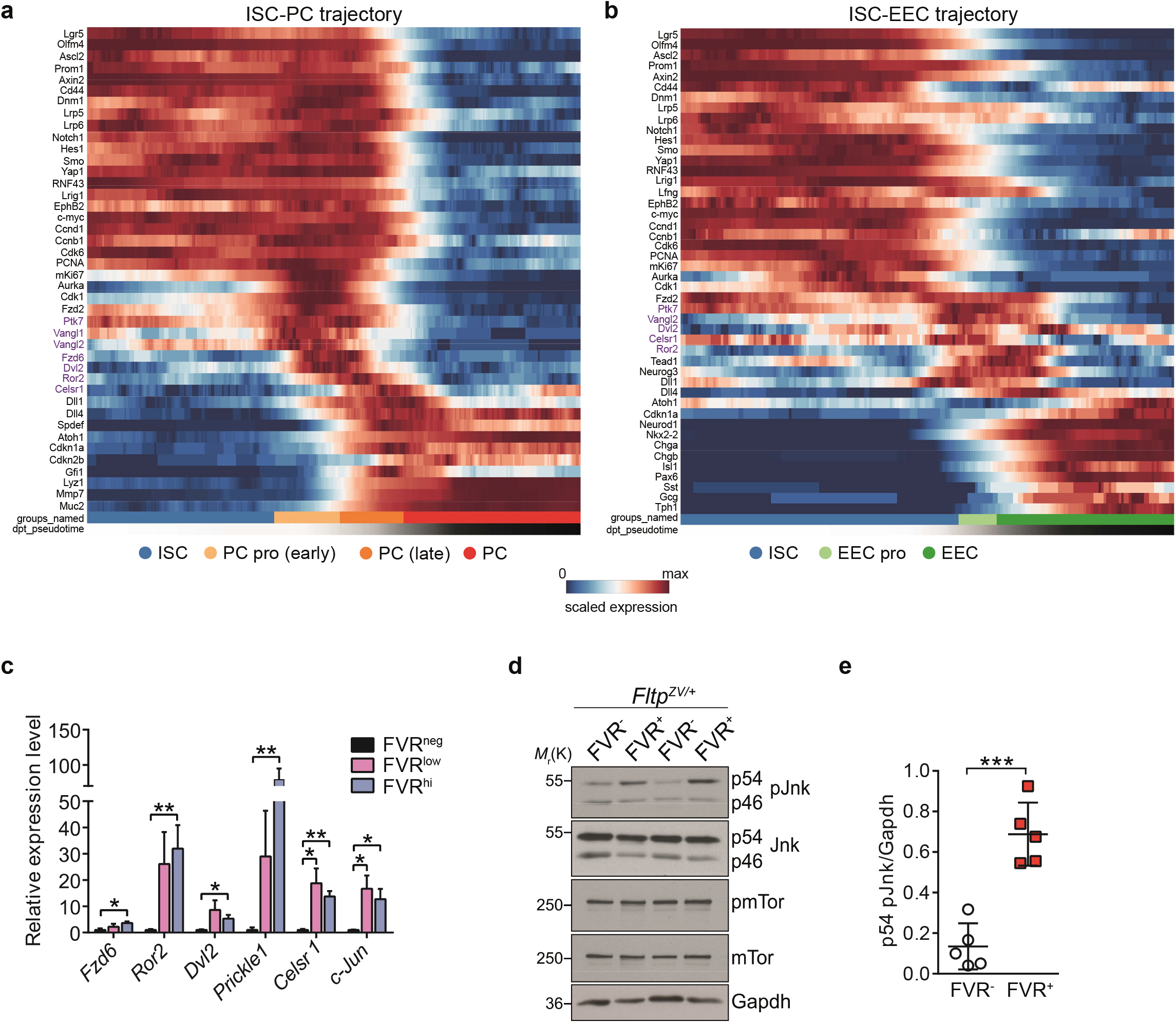
Wnt/PCP signalling is activated during differentiation of ISCs towards PCs and EECs. **a,** Heatmap showing gene expression from scqRT-PCR data along the PC lineage trajectory from ISCs to mature PCs. Cell-type clusters are ordered by the inferred PAGA trajectory and cells by DPT. Expression is shown as the running average over 30 cells scaled to the maximum observed level per gene. Bars at the bottom indicate cell type clusters and DPT. **b,** Heatmap showing gene expression from scqRT-PCR data along the EEC lineage trajectory from ISCs to mature EECs. Cell-type clusters are ordered by the inferred PAGA trajectory and cells by DPT. Expression is shown as the running average over 30 cells scaled to the maximum observed level per gene. Bars at the bottom indicate cell type clusters and DPT. **c,** Expression analysis of Wnt/PCP genes in FVR^hi/low/neg^ crypt cells obtained from *Fltp^ZV/+^* mice by qRT-PCR. *n* (independent experiments) = 3. Error bars represent SEM. Two-tailed Student’s *t*-test, \**P*<0.05, \*\**P*<0.01. **d, e,** Western blot (**d**) and quantification (**e**) of active, phosphorylated (p) Jun N-terminal kinase (Jnk) in flow-sorted FVR^+^ and FVR^-^ crypt cells isolated from adult *Fltp^ZV/+^* reporter mice. Gapdh, mTor and pmTor are presented as controls. *n* (independent experiments) = 5. Error bars represent SD. Two-tailed Student’s *t*-test, \**P*<0.05.

**Extended Data Figure 8.**
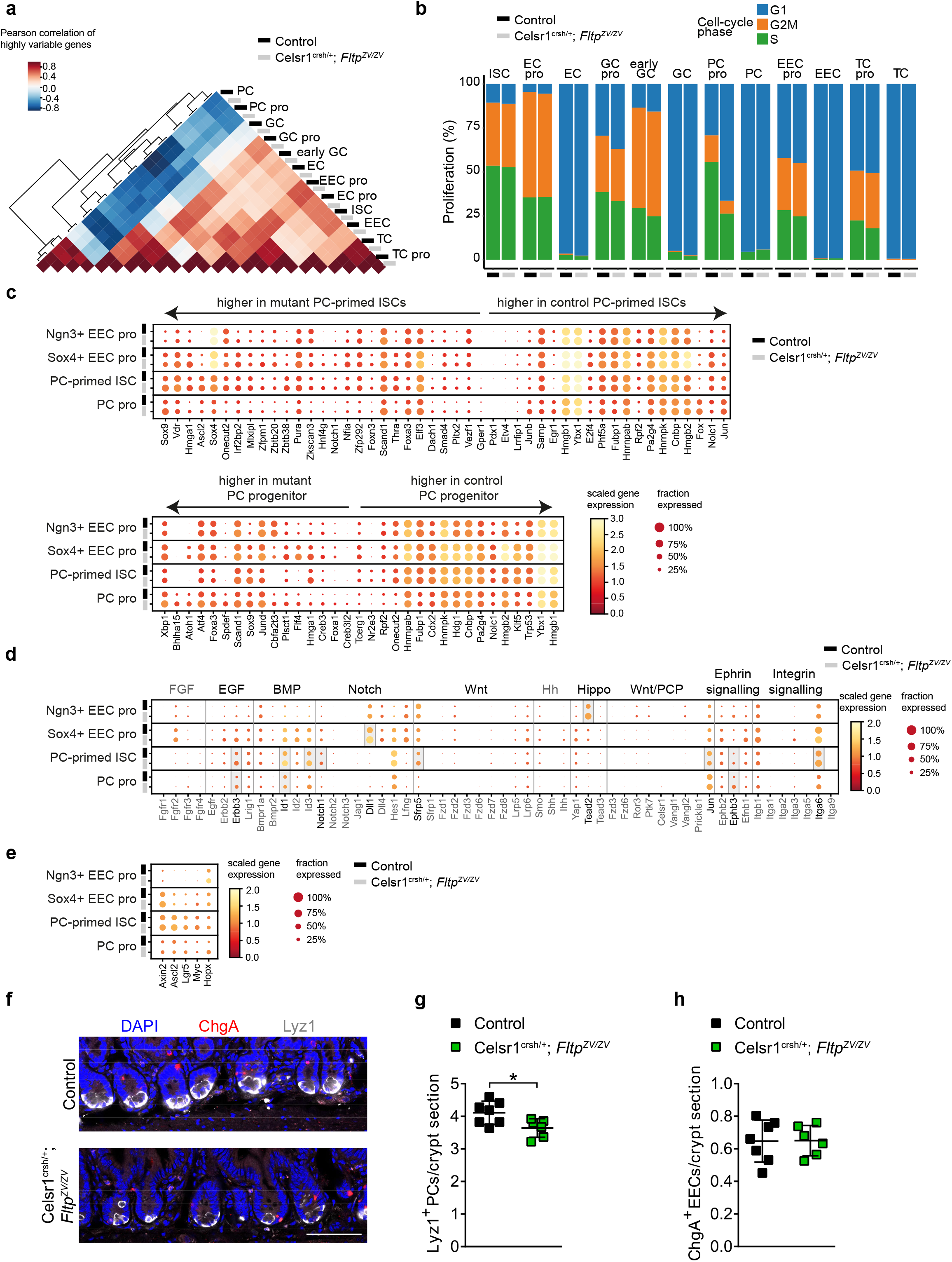
Disturbed Wnt/PCP signalling causes alterations in gene expression pattern and numbers of Paneth cells. **a,** Heatmap showing Pearson correlation of the highly variable genes in all cell types present in Celsr1^crsh/+^; Fltp^ZV/ZV^ mutant and control samples (see Supplementary Table 2). In all cases, Celsr1^crsh/+^; Fltp^ZV/ZV^ and control samples of the respective cell type correlate the most. *n* (mice) = 10 for control mice. *n* (mice) = 4 for Celsr1^crsh/+^; Fltp^ZV/ZV^ compound mutant mice. **b,** Barplot depicting cell-cycle annotation for all cell types into G1 (non-cycling), S and G2M (proliferating) phase. Paneth progenitors in Celsr1^crsh/+^; Fltp^ZV/ZV^ mutant mice are less proliferative than in control mice. All other cell types are not affected in terms of proliferation. *n* (mice) = 4 for control mice. *n* (mice) = 4 for Celsr1^crsh/+^; Fltp^ZV/ZV^ compound mutant mice. **c,** Heatmap depicting differential expression of transcription factors between Celsr1^crsh/+^; Fltp^ZV/ZV^ mutant and control mice and between displayed progenitor groups, ordered by log-fold change in PC-primed ISCs and in PC progenitors. In the EEC progenitors, only *Hmga1* was significantly different in Sox4^+^ early EEC progenitors and none was significantly different in Ngn3^+^ progenitors. Benjamin-Hochberg adjusted p-value < 10^-5^. *n* (mice) = 10 for control mice. *n* (mice) = 4 for Celsr1^crsh/+^; Fltp^ZV/ZV^ compound mutant mice. **d,** Heatmap depicting differential gene expression in specific signalling pathways. Grey boxes in the dotplot highlight significantly different genes. Benjamin-Hochberg adjusted p-value < 10^-5^. *n* (mice) = 10 for control mice. *n* (mice) = 4 for Celsr1^crsh/+^; Fltp^ZV/ZV^ compound mutant mice. **e,** Heatmap depicting expression of Wnt target genes. Expression of *Axin2, Ascl2* and *Lgr5* were significantly different in Celsr1^crsh/+^; Fltp^ZV/ZV^ PC-primed ISCs. Benjamin-Hochberg adjusted p-value < 10^-5^. *n* (mice) = 10 for control mice. *n* (mice) = 4 for Celsr1^crsh/+^; Fltp^ZV/ZV^ compound mutant mice. **f, g, h,** Representative LSM images of duodenal sections stained for ChgA (red, enteroendocrine cells), Lyz1 (white, Paneth cells), DAPI (blue, nucleus) (**f**) and quantification of Paneth cells (**g**) and ChgA^+^ EECs (**h**) in control and Celsr1^crsh/+^; Fltp^ZV/ZV^ compound mutant mice. *n* (mice) = 6 for control mice. *n* (mice) = 7 for Celsr1^crsh/+^; Fltp^ZV/ZV^ compound mutant mice. Error bars represent SEM. Two-tailed Student’s *t*-test, \**P*<0.1. Scale bar, 75 μm.

